# Epigenetic reader ZMYND11 noncanonical function restricts HNRNPA1-mediated stress granule formation and oncogenic activity

**DOI:** 10.1101/2023.11.28.569112

**Authors:** Cheng Lian, Chunyi Zhang, Pan Tian, Qilong Tan, Yu Wei, Qin Zhang, Zixian Wang, Qixiang Zhang, Mengjie Zhong, Li-Quan Zhou, Xisong Ke, Huabing Zhang, Yao Zhu, Zhenfei Li, Jingdong Cheng, Gong-Hong Wei

## Abstract

ZMYND11 encodes an epigenetic reader of histone methylation, functioning as a transcriptional corepressor. However, whether and how ZMYND11 contributes to cancer progression and therapy remains unclear. Here we report that ZMYND11 downregulation is prevalent in cancers and profoundly correlates with adverse prostate cancer patient outcomes. Depletion of ZMYND11 promotes tumor cell growth, migration and invasion *in vitro* as well as tumor formation and metastasis *in vivo*. Mechanistically, we find that ZMYND11 exhibits tumor suppressive roles through recognizing arginine-194-methylated HNRNPA1 dependent on its MYND domain, thereby squeezing HNRNPA1 in nucleus and inhibiting the formation of stress granules in cytoplasm. Furthermore, ZMYND11 antagonizes HNRNPA1-driven high PKM2/PKM1 ratio and counteracts PKM2-induced aggressive tumor phenotype. Remarkably, ZMYND11 recognition of HNRNPA1 could be disrupted by pharmaceutical inhibition of arginine methyltransferase PRMT5 while ZMYND11 low-expressing tumors are sensitive to the treatment of PRMT5 inhibitors. Collectively, our study unravels a novel and noncanonical function of ZMYND11 as the nonhistone methyl reader and discovers a mechanism for the requirement of arginine-methylation-mediated ZMYND11-HNRNPA1 association to restrict tumor progression and offers cancer therapeutic targets and potential biomarkers.

## Introduction

Cancer is a complex disease in which oncogene and tumor suppressor interplays are fundamental basis of both tumor progression and therapeutic response ^1^. *ZMYND11* (also known as BS69) encodes a candidate tumor suppressor consisting of a plant homeodomain (PHD), a bromodomain, a Pro-Trp-Trp-Pro (PWWP) domain, and a myeloid, Nervy, and DEAF-1 (MYND) domain. Previous studies have demonstrated that ZMYND11 is an epigenetic reader specific for H3.3K36me3 recognition via the PHD-bromo-PWWP (PBP) domain ^2, 3^. Moreover, ZMYND11 functions as a transcriptional corepressor by modulating RNA polymerase II (Pol II) at the elongation stage ^2, 3^. Known data showed that the MYND domain at C-terminus of ZMYND11 could mediate protein-protein interaction with certain transcription factors and splicing-related proteins ^3, 4^, contributing to transcriptional repression of downstream oncogenes ^5, 6^. These findings suggest diverse functions of ZMYND11 in reading histone modifications and transcriptional regulation via different cooperative proteins. While it is clear how ZMYND11 is involved in recognition of H3.3K36me3, it remains unclear whether ZMYND11 possesses unconventional functions and antagonize any critical oncoprotein, thereby suppressing cancer progression.

Heterogeneous nuclear ribonucleoprotein A1 (HNRNPA1) has been reported to play fundamental roles in gene expression and functions as an RNA-binding protein (RBP) harboring two RNA recognition motifs (RRM) and a glycine-rich domain including several Arg–Gly–Gly (RGG) tripeptide repeats which could mediate cellular compartmentalization, protein-protein interaction, and RNA-binding ^7–9^. Previous studies showed that HNRNPA1 was involved in the regulation of RNA Pol II associated nascent transcripts and thus RNA splicing of critical genes such as the pyruvate kinase gene (PKM). Specifically, HNRNPA1 promotes the excision of exon 9 of the PKM pre-mRNA to selectively induce oncogenic PKM2 isoform, a known critical determinant of the Warburg effect, thereby promoting aerobic glycolysis and favoring cancer cell growth and survival ^7–9^. Moreover, accumulating evidence demonstrated that HNRNPA1 was also involved in stress granule assembling into liquid-liquid phase separation ^10^, implicating as a driver force in a variety of human disorders including cancers. The formation of stress granules endows malignant cells with survival advantages and chemotherapy resistance ^11, 12^. Targeted stress granules assembly may represent a potential therapeutic strategy to overcome primary and acquired chemotherapy resistance and improve efficacy ^13–16^. Nevertheless, the molecular mechanisms governing oncogenic function of HNRNPA1 in cancers remain elusive.

In this study, we report that ZMYND11 plays essential roles in suppressing tumor cell proliferation, metabolism and cancer progression through a noncanonical function in binding with the PRMT5-mediated arginine methylation of HNRNPA1, thereby antagonizing cancer driver capacity of HNRNPA1. Furthermore, we show that ZMYND11 underexpression tumors indicate a sensitive therapeutic response to the type II arginine methyltransferase PRMT5 inhibitors.

## Result

### ZMYND11 is profoundly downregulated in cancers and its downregulation correlates with adverse events and poor PCa patient outcomes

To gain a broad insight into dysregulation of the epigenetic reader ZMYND11 in cancers, we initially examined the expression of ZMYND11 across various types of human tumors and adjacent normal tissue samples. The results demonstrated an apparent and frequent downregulation of ZMYND11 in non-epithelial cancers (brain, mesothelioma) and multiple epithelial cancer types (cervix, colorectal, esophagus, stomach, pancreas, prostate) especially significant in PCa **(Figure 1A)**. Therefore, this observation motivated us for subsequent study of the clinical relevance and consequences of ZMYND11 downregulation in PCa, the most prevalent and lethal urological cancer in men. We subsequently confirmed downregulation of ZMYND11 in The Cancer Genome Atlas Prostate Adenocarcinoma (TCGA PRAD) data **(Figure S1A)**. Notably, ZMYND11 showed consistent downregulation in metastatic PCa samples from several independent human PCa clinical datasets ^17–20^ **(Figures 1B and S1B)**, and its downregulation markedly correlated with high Gleason score, advanced tumor stage and elevated prostate-specific antigen (PSA) levels ^21–23^ **(Figures 1C and S1C)**. To further define the importance of ZMYND11 in PCa, we compared ZMYND11 expression at both mRNA and protein levels in prostate tissue lysates derived from mice harboring *Pbi–Cre*–mediated deletion of *Pten (Pten^-/-^)* and having indolent PCa. As expected, we observed marked underexpression of ZMYND11 in murine PCa tumors than normal prostate glands **(Figures 1D and S1D).** To extend the observation, we performed immunohistochemistry (IHC) staining of ZMYND11 in two independent tumor tissue microarrays (TMA) with an antibody against ZMYND11. The results showed that ZMYND11 protein levels were obviously decreased in PCa patient tumor samples compared with surrounding non-tumor tissues **(Figures 1E and 1F).** Collectively, these findings suggest a potential role of ZMYND11 in restricting PCa progression to advanced stage and metastasis.

**Figure 1:**
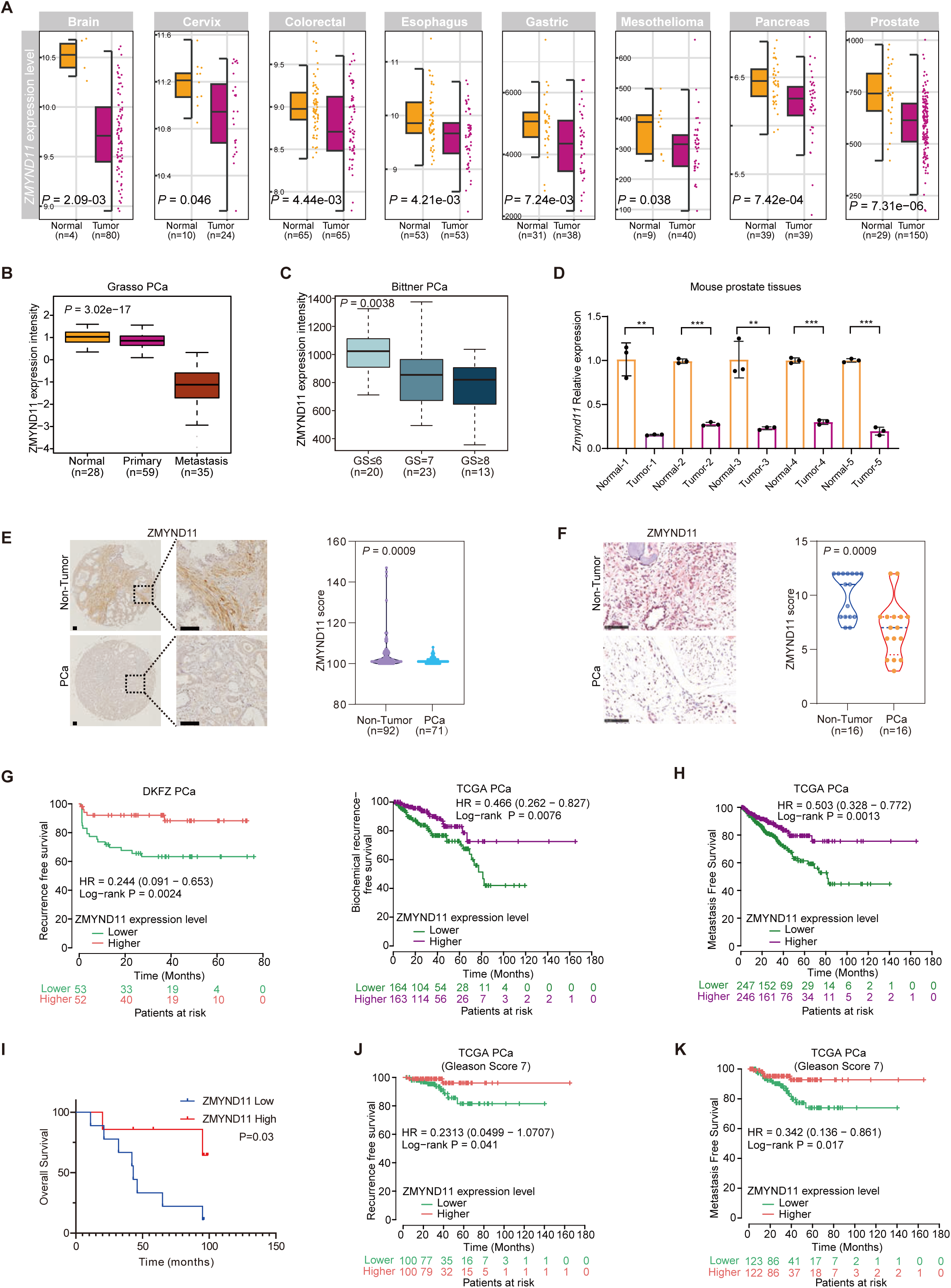
ZMYND11 is profoundly downregulated in cancers and its downregulation correlates with adverse events and poor PCa patient outcomes. **(A)**, Downregulation of *ZMYND11* gene expression in several types of cancers (red) compared to healthy tissues (yellow). *P* values were assessed by the Wilcoxon test while effect sizes were estimated using Cohen’s distance: *d*_brain_=1.96, *d*_cervix_=0.86, *d*_colon_=0.56, *d*_esophagus_=0.64, *d*_gastric_=0.7, *d*_mesothelioma_=0.64, *d*_pancreas_=0.77, *d*_prostate_=0.99. **(B)**, ZMYND11 mRNA expression was significantly downregulated in human metastatic prostate tumors. **(C)**, Association of ZMYND11 downregulation with high Gleason grade cancer. *P* values examined by the Kruskal-Wallis test in (**B**)-(**C**). **(D)**, Expression of *Zmynd11* in five paired PCa specimens and adjacent normal tissues of murine prostate. **(E)**, Protein expression levels of ZMYND11 in paraffinized prostate tumor tissue microarrays (TMA) of Tongji PCa cohort determined by immunostaining. Original magnification, 100×; insets, 400x; Scale bar, 100 μm. **(F)**, ZMYND11 protein expression were determined in Fudan cohort of PCa using immunohistochemistry. The IHC scores calculated with staining areas (see Methods). Original magnification, 400×; Scale bar, 100 μm. **(G)**, Kaplan-Meier analysis of biochemical recurrence-free survival in prostate tumors with high or low levels of ZMYND11 in two independent cohorts of PCa. **(H)**, Metastasis free survival analysis of 493 PCa patient with tumors expressing high or low mRNA levels of ZMYND11. **(I)**, Kaplan-Meier overall-survival analysis in a TMA cohort of patients with PCa tumors having higher protein expression levels of ZMYND11 (top 50%; n = 35) or lower (bottom 50%; n =35). **(J and K)**, Lower expression levels of ZMYND11 indicates predictive values for recurrence (**J**) and metastasis-free (**K**) survival in patient group with Gleason score 7 (intermediate-risk PCa). In **D**-**F**, statistical significance assessed using the two-tailed Student’s t test. **p < 0.01, ***p < 0.001. In **G**-**K**, *p* values examined by the log-rank test.

To further address the clinical relevance of these observations, we performed prognosis correlation analysis across multiple independent cohorts of PCa patients with long-term clinical outcomes. We found that the patient group with lower mRNA expression of ZMYND11 had increased risk for postoperative biochemical relapse and tumor progression to metastasis as well as shorter time for overall survival ^18, 21, 23^ **(Figures 1G,1H and S1E,S1F).** In line with these observations, a Kaplan-Meier analysis of our TMA immunostaining results revealed that the PCa patients with lower protein levels of ZMYND11 displayed significantly shortened time for overall survival compared with the patients with higher ZMYND11 expression levels **(Figure 1I).** Given that ZMYND11 expression status was reproducibly found to be associated with PCa prognosis, we next asked whether ZMYND11 levels shows any predictive values for low- and high-risk PCa. Indeed, we found that ZMYND11 mRNA levels are obviously predictable for biochemical recurrence and metastasis of the patients with Gleason score 7 (intermediate risk; **Figures 1J,1K and S1G),** but not for the patients groups with Gleason score ≤6 (low risk; **Figure S1H)** or ≥8 (high risk; **Figure S1I),** indicating ZMYND11 as a potential prognostic marker to stratify the intermediate risk PCa patients who may recur or progress to metastasis. Taken together, these data demonstrate that downregulation of ZMYND11 frequently occurs during PCa tumor progression to advanced stage and correlates with unfavorable prognosis in PCa patients, suggesting that ZMYND11 could act as a tumor suppressor and dampened ZMYND11 may contribute to PCa tumorigenesis and metastasis.

### Dampened ZMYND11 expression promotes PCa cell proliferation and metastasis *in vitro* and *in vivo*

We next sought to investigate the effect of ZMYND11 on PCa tumor cellular phenotypes and invasiveness and thus conducted knockdown assays using lentivirus-mediated short hairpin RNA (shRNA) or synthetic small interfering RNA (siRNA) against ZMYND11 in several PCa cell models **(Figures 2A and S2A)**. The results consistently showed that dampened ZMYND11 greatly stimulated colony formation, cell growth, migration, and invasion of the PCa cell lines 22Rv1, DU145, and LNCaP **(Figures 2B-2E and S2B-S2E)**. We further examined the effect of ZMYND11 knockdown on tumor growth *in vivo* in the xenograft mouse model. The results showed that the growth of xenograft PCa tumors composed of 22Rv1 cells with stable silencing of ZMYND11 were significantly increased compared with xenografts composed of control cells **(Figure 2F)**. Furthermore, we established the 22Rv1 cell models stably expressing firefly luciferase for metastatic tumor growth assays and quantitated the lung metastatic mouse model after tail vein injections by bioluminescent imaging (BLI). Our weekly BLI analysis showed that ZMYND11 knockdown greatly promoted lung metastasis **(Figure 2G)**. We subsequently performed FACS separation of the GFP-positive circulating cancer cells from the blood and observed significantly higher amount of circulating 22Rv1 cells with ZMYND11 knockdown than controls **(Figure 2H)**, indicating that the cells with dampened ZMYND11 may possess enhanced survival capacity in the blood stream and thereby facilitating metastasis to distant organs. Altogether, these data have established the importance of ZMYND11 downregulation for PCa cell growth and metastasis *in vitro* and *in vivo* and are consistent with the above observed link between low ZMYND11 levels and prostate tumor aggressive phenotype as well as PCa progression and poor prognosis in the clinical setting.

**Figure 2:**
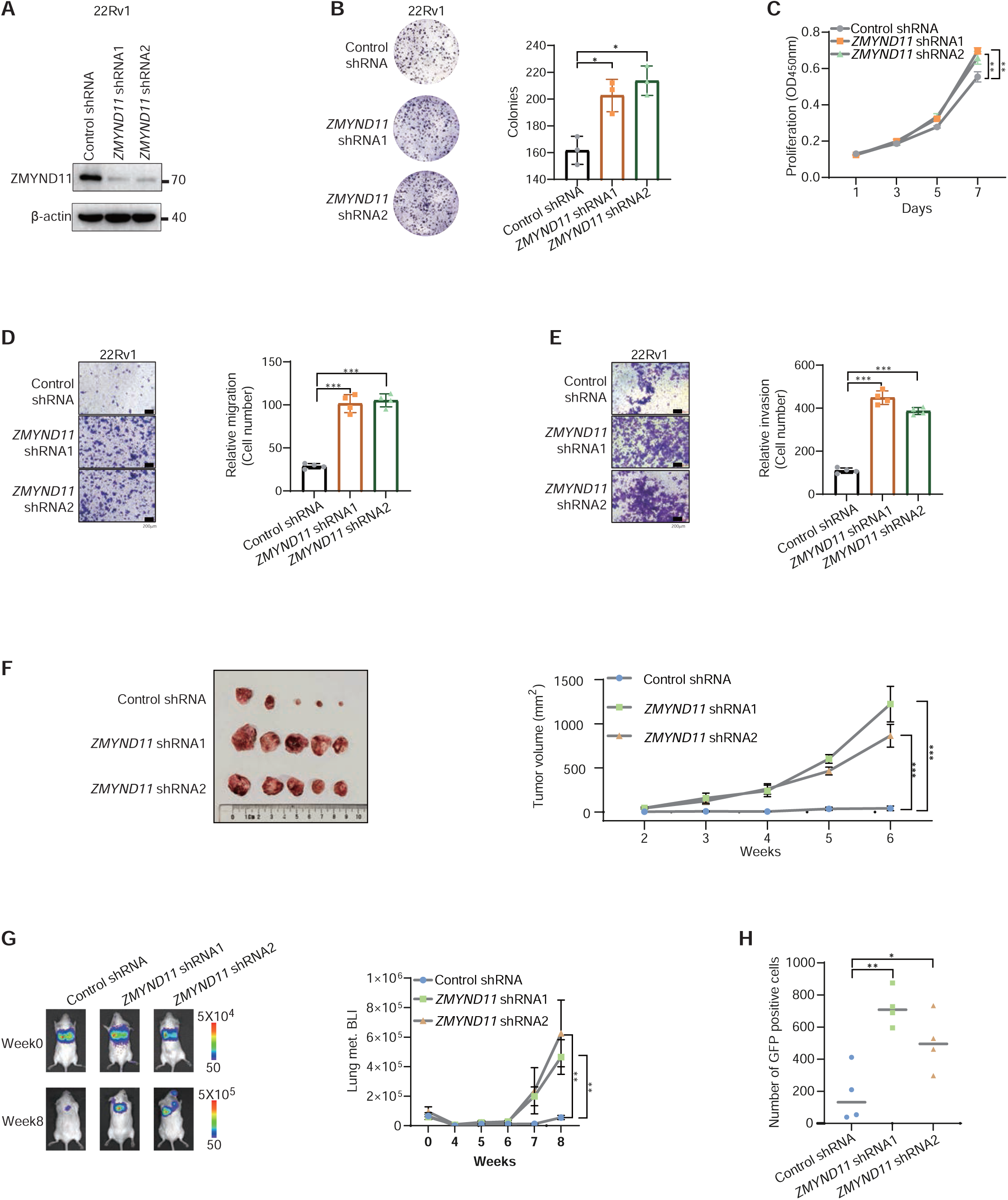
Dampened ZMYND11 expression promotes PCa cell proliferation and metastasis *in vitro* and *in vivo*. **(A)** Efficiency of shRNA-mediated ZMYND11 knockdown was determined by immunoblotting. **(B)** Colony-forming units of 22Rv1 cells stably expressing control or shRNAs against ZMYND11. **(C)** Proliferation rates of 22Rv1 cells measured at indicated times (absorbance at 450 nm; mean ± SD of three independent experiments). **(D and E)**, Representative images and quantification of relative migration **(D)** and invasion **(E)** for the cells stably expressing the indicated shRNAs. **(F)** Representative images of the tumor xenografts harvested 6 weeks after subcutaneous injection of 1 x 10^7^ 22Rv1 cells with the indicated control or shRNA-mediated stable silencing of ZMYND11 into the nude mouse (left). Growth curves of 22Rv1 xenografts at the indicated time points (right). **(G)** NOD-SCID mice treated with tail vein injection of 22Rv1 cells having stable knockdown of ZMYND11 and stable transfection of the firefly luciferase. Representative images at the experimental endpoint and weekly quantitation of bioluminescent imaging (BLI) analyses of mouse lung metastasis. **(H)** Fluorescence activated cell sorting (FACS) analysis of the circulating GFP-positive 22Rv1 cell variants (CTCs) in the blood of SCID mice. The scatter plot shows the number of CTCs recovered from each mouse injected with control or shRNA-mediated ZMYND11 knockdown cells. In **B**-**H**, data shown are mean ± SD Error bars, *p < 0.05, **p < 0.01, ***p < 0.001, two-tailed Student’s t-test.

### ZMYND11 interacts with HNRNPA1 protein by MYND domain and inhibits HNRNPA1-mediated the formation of stress granule

To gain insights into the molecular mechanisms underlying the role of ZMYND11 in regulating cancer cell growth and metastasis, we sought to elucidate new interacting proteins of ZMYND11. Thus, we performed immunoprecipitation experiment in the PCa LNCaP cells with an antibody against endogenous ZMYND11 followed by mass spectrometry (MS) analysis of the precipitated proteins. A total of 37 proteins were robustly identified to be associated with ZMYND11 in the three independent experiments **(Figure 3A)**. Gene Ontology (GO) analysis indicated that most proteins associated with ZMYND11 are enriched in the functional category of RNA splicing **(Figure S3A)**, consistent with the previous observation that ZMYND11 regulates RNA alternative splicing (AS) ^2, 3^. While ranking the candidate ZMYND11 interaction proteins, HNRNPA2B1 and HNRNPA1 are the two top hits among three repetitions of protein complex identification **(Figure 3B)**. Co-immunoprecipitation (Co-IP) assays further demonstrated HNRNPA1, but not HNRNPA2B1, could interact with ZMYND11 in transfected cells **(Figures 3C and S3B)**. The physiological interaction between ZMYND11 and HNRNPA1 was confirmed at the endogenous levels in 22Rv1 cells **(Figure 3D)** and by an immunofluorescence assay, showing that endogenous ZMYND11 colocalizes with HNRNPA1 in the nucleus **(Figure 3E)**. ZMYND11 could also interact with HNRNPA1 in the lung cancer cell line A549 and the breast cancer cell line MDA-MB-231 **(Figure S3C)**, suggesting that ZMYND11 binding to HNRNPA1 may be prevalent in other types of cancers.

**Figure 3:**
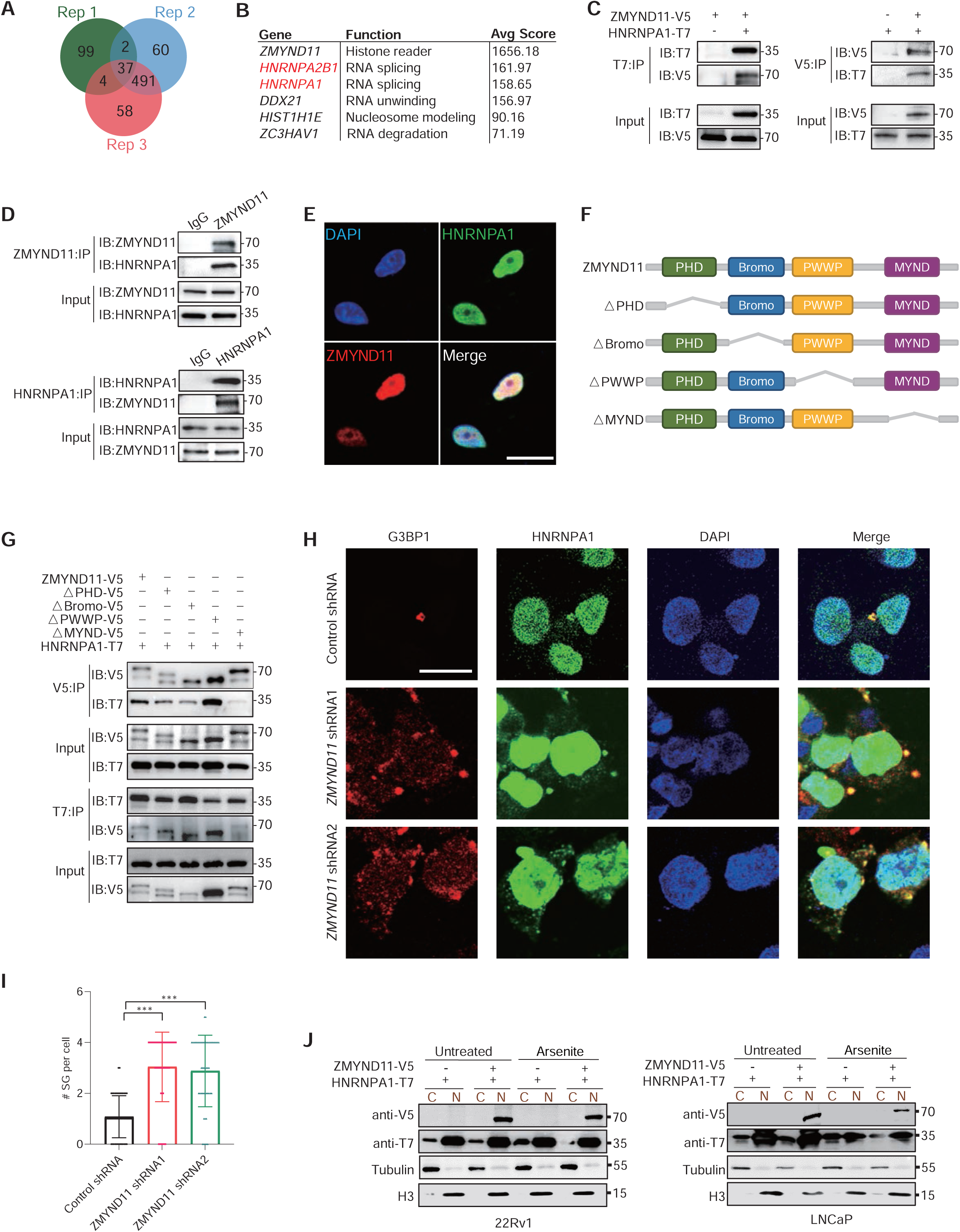
ZMYND11 physically interacts with HNRNPA1 *in vitro* and *in vivo* and inhibits HNRNPA1-mediated formation of stress granules. **(A)** Venn diagram showing the overlap between three replicates of the identified ZMYND11-interaction proteins in LNCaP cells using co-immunoprecipitation coupled to mass spectrometry (Co-IP/MS). **(B)** Table showing top-scoring ZMYND11-interaction proteins reproducibly identified by Co-IP/MS as indicated in **A**. **(C)** Immunoprecipitation (IP)-Western blot for analyzing the interaction between ZMYND11 and HNRNPA1 in HEK293T cells cotransfected with the indicated V5- or T7-tagged plasmids. IB, immunoblot. **(D)** Immunoprecipitation-Western blot analysis of an endogenous interaction of ZMYND11 with HNRNPA1 in 22Rv1 cells. IgG, immunoglobulin. IB, immunoblot. **(E)** Immunofluorescence (IF) staining and colocalization analysis of ZMYND11 and HNRNPA1 in 22Rv1 cells. Scale bar, 25 μm. **(F)** Schematic illustration of the full-length ZMYND11 and the indicated domain deletion mutants. **(G)** Immunoprecipitation-Western blot analysis of the interactions between wild type ZMYND11 or its mutants and HNRNPA1. Crude total cell lysates were extracted from HEK293T cells co-expressing T7-tagged HNRNPA1 and the indicated V5-tagged ZMYND11 variants. **(H)** 22Rv1 cells stably expressing control or shRNAs against ZMYND11, treated with sodium arsenite (SA, 0.5mM) for 1h followed by immunostaining analysis of stress granule (SG) formation. Scale bar, 25 μm. **(I)** The number of SGs per cell and ZMYND11 or HNRNPA1 colocalized SGs each cell were quantified. **p < 0.01, ***p < 0.001 evaluated by two-tailed Student’s t-test. **(J)** Immunoblotting analysis of ZMYND11 or HNRNPA1 in the nuclear and cytoplasmic fractions of the tested PCa cell lines 22Rv1 and LNCaP, respectively.

We next assessed which domain of ZMYND11 protein mediated the interaction with HNRNPA1. Given that ZMYND11 contains plant homeodomain (PHD), bromodomain (Bromo), Pro-Trp-Trp-Pro (PWWP), and myeloid, Nervy, and DEAF-1 (MYND) domains, we generated four domain-deletion mutant constructs with a C-terminal V5 tag (**Figure 3F**). Co-IP experiments demonstrated that only deletion of the MYND domain, but not PWWP, Bromo, or PHD domain of ZMYND11, greatly diminished full-length ZMYND11 binding to HNRNPA1, indicating that the MYND domain of ZMYND11 is essential for its interaction with HNRNPA1 **(Figure 3G)**. These data indicated that ZMYND11 interacts with HNRNPA1 protein by MYND domain.

Both *in vitro* and *in vivo* evidence as investigated above support a direct interaction between tumor suppressor ZMYND11 and oncogenic HNRNPA1 ^8, 24, 25^, we thus sought to explore potential functional consequence of this novel protein association. Notably, it has been shown that HNRNPA1 was involved in stress granules assembly into liquid-like droplets called liquid-liquid phase separation while stress granule formation could play roles in preventing cancer cell apoptosis, tumor progression and chemotherapeutic resistance ^14, 15, 26, 27^. We hence investigated the role of ZMYND11-HNRNPA1 association in stress granule and stressed the cells with sodium arsenite (SA; 0.5mM, 1h). Immunofluorescence analysis revealed that the cytoplasm localized HNRNPA1 was increased in the cells eliciting a constitutively active form of stress granules that colocalized with the stress granule marker G3BP1 ^28^, whereas ZMYND11 as a control cannot assemble stress granules in response to SA treatment **(Figures S3D-S3E)**. We detected the endogenous HNRNPA1 under SA treatment. Notably, the knockdown of ZMYND11 showed increased stress granules formation that co-localized with HNRNPA1 **(Figures 3H-3I)**. When there was simultaneous ectopic expression of ZMYND11 and HNRNPA1 in the cells, the stress-induced cytoplasmic accumulation of HNRNPA1 and stress granule formation of HNRNPA1 both were substantially inhibited **(Figures S3F-S3G)**. In line with this, the nuclear and cytoplasmic separation experiments showed that the cytoplasmic HNRNPA1 levels decreased and this was likely due to ZMYND11 physical association with and thereby sequestering HNRNPA1 in the nucleus under the stress treatment **(Figure 3J)**. Taken together, these findings indicate that ZMYND11 interacts with HNRNPA1 protein by the MYND domain and could restrict cytoplasmic translocation of HNRNPA1 to the stress granules in the cells undergoing stress.

### ZMYND11 antagonizes the oncogenic HNRNPA1-induced aggressive cellular phenotype of PCa depending on the MYND domain

HNRNPA1 was widely studied in different types of human cancers while few were reported in PCa ^24, 25, 29^. We therefore next assessed the influence of HNRNPA1 on tumor cellular phenotypes in PCa and demonstrated that shRNA or siRNA-mediated silencing of HNRNPA1 led to suppressed cell growth, colony formation, migration, and invasion in PCa cell lines 22Rv1, DU145, and LNCaP **(Figures 4A-4E and S4A-S4E)**. Growing evidence illustrated that generally, HNRNPA1 acted as an oncogene and its upregulation was associated with higher pathological stages and poor prognosis in multiple types of cancers ^29^. To examine whether this is the case in PCa, we assessed HNRNPA1 mRNA levels across multiple independent clinical datasets. The results showed that HNRNPA1 expression was dramatically upregulated during PCa tumor progression to advanced/metastatic stage in four cohorts of datasets ^18, 20, 30, 31^ **(Figures 4F and S4F)**. In addition, HNRNPA1 expression was increased in patients with high Gleason score while increased expression of HNRNPA1 correlated with elevated risk of disease relapse and metastasis ^20, 23^ **(Figurse 4G and S4G)**. We also measured the protein levels of HNRNPA1 in PCa tumor from mice with prostate-specific deletion of *Pten (Pbi–cre; Pten^loxP/loxP^)* and normal murine prostate gland, demonstrating that HNRNPA1 was increased in tumor tissues **(Figure S4H)**. To expand the observation on clinical relevance of HNRNPA1 in PCa, we measured HNRNPA1 protein levels in our collection of PCa patient samples and compared with surrounding non-tumor tissues by IHC. The results demonstrated that HNRNPA1 expression was increased in tumor specimens **(Figures 4H and 4I)** and importantly, the patients with higher HNRNPA1 levels exhibited much shorter overall survival than those with lower HNRNPA1 expression **(Figure 4J)**. Further analysis of several public datasets revealed similar results to the observations in our cohorts, showing that the time to disease relapse and metastasis was substantially shorter in the group of PCa patients with higher levels of HNRNPA1 ^23, 32^ **(Figures 4K and S4I)**. Notably, we found that HNRNPA1 expression is clearly predictable for disease recurrence of the PCa patients with Gleason score 7 but not for the patients with Gleason score ≤6 or ≥8 (intermediate risk; **Figures 4l and S4J**), suggesting a prognostic potential of HNRNPA1 oncogene in patient stratification.

**Figure 4:**
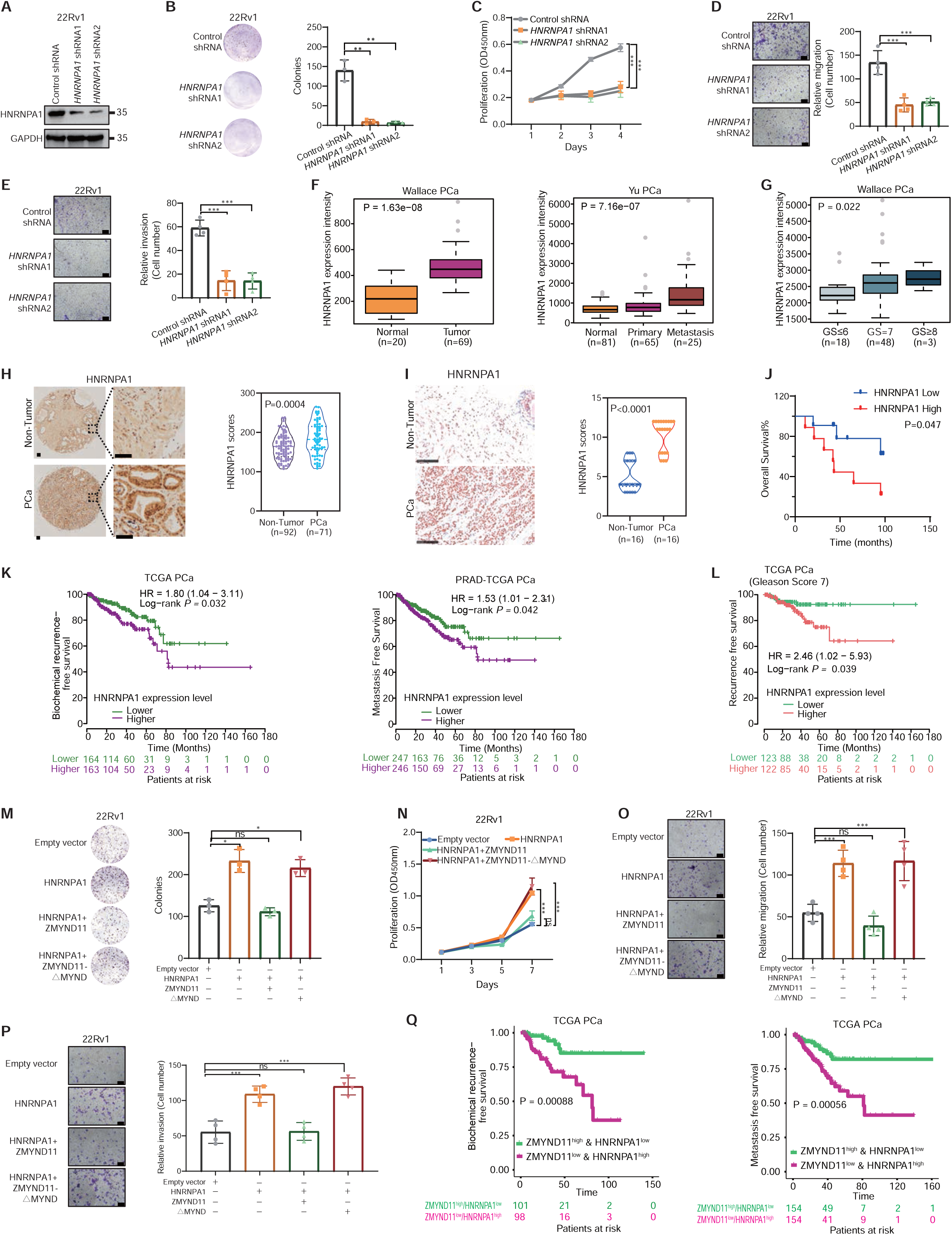
ZMYND11 antagonizes oncogenic function of HNRNPA1 in PCa dependent on its MYND domain. **(A)** Knockdown efficiency of shRNAs against HNRNPA1 determined by western blot analysis. **(B)** Colony-forming units of 22Rv1 cells stably expressing control or shRNAs against HNRNPA1. **(C)** Proliferation capacity of 22Rv1 cells measured at the indicated times. Absorbance at 450 nm; mean ± SD of three independent experiments. **(D and E)**, Representative images and quantification of relative migration **(D)** and invasion **(E)** for the cells stably expressing the indicated shRNAs. Error bars, ± SD of triplicate experiments. **(F)** HNRNPA1 expression levels are significantly upregulated in human metastatic prostate tumors. *P* values determined by the Wilcoxon and Kruskal-Wallis tests, respectively. **(G)** HNRNPA1 upregulation correlates with higher Gleason grade cancer. *P* values assessed by the Kruskal-Wallis test. **(H)** Immunohistochemistry on the expression of HNRNPA1 and evaluation of HNRNPA1 staining intensity in a paraffinized prostate tissue microarray consisting of 163 samples. Original magnification, 100×; insets, 400×; Scale bar, 100 μm. **(I)** Immunostaining for HNRNPA1 was performed on another cohort of prostate specimens. Representative images and quantification of HNRNPA1 protein staining are shown. Original magnification, 400×; Scale bar, 100 μm. **(J)** Kaplan-Meier analysis of overall survival in the Tongji cohort of patients with PCa. Patients were stratified into two groups according to the state of HNRNPA1 expression, higher (top 50%; n = 35) or lower (bottom 50%; n =35). **(K)** Kaplan-Meier analysis showed that HNRNPA1 upregulation is significantly associated with elevated risk of biochemical recurrence or metastasis in PCa subsets within the TCGA database. **(L)** High expression levels of HNRNPA1 indicates predictive values for recurrence-free survival in patient group with Gleason sum score 7 PCa (intermediate-risk). **(M-P)**, Representative images and quantification of colony formation (**M**), cell proliferation (**N**), migration (**O**), and invasion (**P**) for 22Rv1 cells transfected with empty vector, HNRNPA1 alone, or together with the indicated full-length ZMYND11 or MYND-domain deletion mutant. Scale bar, 200 μm. **(Q)** Kaplan-Meier analysis of biochemical recurrence or metastasis-free survival in the TCGA cohort of PCa patients stratified into two groups with ZMYND11^low^/HNRNPA1^high^ and ZMYND11^high^/HNRNPA1^low^ expression, respectively. Error bars, ±SD (**B**-**E**, **M**-**P**). ns, not significant, *p < 0.05, **p < 0.01, ***p < 0.001, two-tailed Student’s t tests.

We next examined the effects of ZMYND11 on HNRNPA1 function in prostate oncogenesis and thus performed forced co-expression of full-length or MYND-domain-deleted ZMYND11 with HNRNPA1 in PCa cells. The results demonstrated that full-length ZMYND11, but not MYND-domain-deleted variant antagonized enhancement of cell growth, colony formation, migration, and invasion induced by HNRNPA1 overexpression **(Figures 4M-4P and S4K-S4N)**, suggesting that ZMYND11 compromised oncogenic function of HNRNPA1 depending on their physical interaction mediated by MYND domain. Given that ZMYND11 downregulation or HNRNPA1 upregulation correlate with adverse outcomes of PCa patients, we next addressed whether the observations have relevant clinical impacts. As indicated in **Figure 4Q**, patients with ZMYND11^low^/HNRNPA1^high^ expression demonstrated tremendously shorter biochemical recurrence- or metastasis-free survival than the ones with ZMYND11^high^/HNRNPA1^low^ expression, suggesting a synergistic prognostic effect of both genes on PCa severity. Taken together, our data demonstrate oncogenic and clinical impacts of HNRNPA1 on PCa progression and that ZMYND11 attenuates oncogenic capabilities of HNRNPA1 through MYND domain of ZMYND11 dependent binding to HNRNPA1.

### ZMYND11 regulates HNRNPA1-mediated alternative splicing of *PKM* and counteracts PKM2-induced aggressive phenotype of prostate cancer cells

HNRNPA1 was reported to promote the excision of exon 9 of the PKM pre-mRNA, thereby selectively inducing oncogenic PKM2 isoform formation, a known critical determinant of the Warburg effect that promotes aerobic glycolysis and cancer cell survival ^7–9^ **(Figure S5A)**. Here, we found that ZMYND11 could physically interact with HNRNPA1 via MYND domain, and this interaction was essential for ZMYND11 to antagonize the HNRNPA1-induced aggressive phenotypes of prostate cancer cells (**Figures 3-4 and S3-S4**); hence, we next sought to determine whether ZMYND11 regulated alternative splicing of PKM pre-mRNA and impacted the function of cancer cells. Notably, the full-length ZMYND11, but not MYND-domain-deleted ZMYND11 inhibited PKM2 and promoted PKM1 isoform formation in both mRNA **(Figure 5A)** and protein level **(Figure 5B)**. In contrast, ZMYND11 knockdown by shRNA reduced PKM1 and enhanced PKM2 isoform formation **Figures 5C and 5D)**; consistently, the full-length ZMYND11 blocked the stimulatory effects of HNRNPA1 on PKM2 formation **(Figures 5E and 5F)**, further indicating the inhibitory effect of ZMYND11 on HNRNPA1-mediated PKM splicing. PKM2 is a splicing form of pyruvate kinase, as a vital determinant of cancer metabolism phenotype and provides selectively proliferative capability for tumor cell growth *in vivo* ^33, 34^. We therefore sought to explore whether ZMYND11 counteracted the PKM2 overexpression-induced aggressive phenotype of the PCa cells. The results showed that the full-length ZMYND11 but not the MYND-domain-deleted variant antagonized enhancement of cell growth, migration, and invasion induced by PKM2 overexpression **(Figures 5G-5J and S5B-S5E)**.

**Figure 5:**
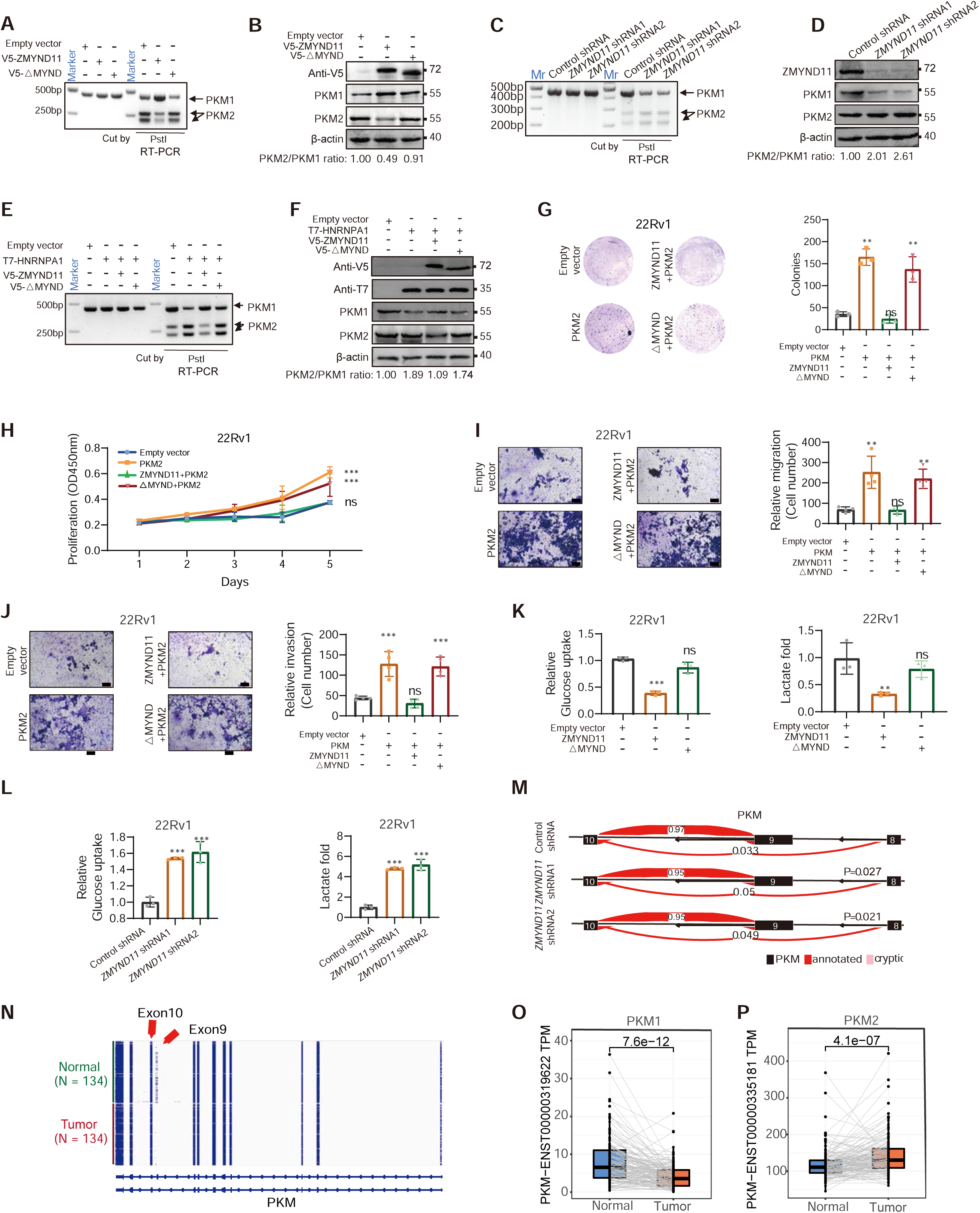
ZMYND11 regulates HNRNPA1-mediated alternative splicing of PKM and counteracts PKM2-induced aggressive cellular phenotype in PCa. **(A-D)**, RNA (**A, C**) or protein (**B, D**) was extracted from 22Rv1 cells either transfected with full-length ZMYND11 or the indicated mutant (**A, B**) or stably expressing the ZMYND11-targeting shRNAs (**C, D**) followed by the examination of PKM isoforms both at the mRNA (**A, C**) and protein levels (**B, D**). **(E and F)** 22Rv1 cells transfected with HNRNPA1 alone or together with either full-length ZMYND11 or the MYND-domain-deletion mutant were assayed for PKM splicing at the mRNA (**E**) and protein levels (**F**). **(G-J)**, Representative images and quantification of colony forming (**G**), cell proliferation (**H**), migration (**I**), and invasion (**J**) in 22Rv1 cells co-transfected with PKM2 alone or together with either the full-length ZMYND11 or the MYND-domain-deletion mutant. Mean ± SD of triplicate experiments. *p < 0.05, **p < 0.01, ***p < 0.001, Student’s t test. Scale bar, 200 μm. **(K and L)** Relative glucose uptake and lactate production in 22Rv1 cells overexpressing ZMYND11 or its MYND-domain-deletion mutant (**L**) and stably expressing ZMYND11 shRNA (**L**) compared to control cells. Error bars, ± SD (**K**-**L**), n = 3 biological replicates. *p < 0.05, **p < 0.01, ***p < 0.001, Student’s t tests. **(M)** Visualization of PKM splicing using LeafViz examined by RNA sequencing in 22Rv1 cells stably expressing either ZMYND11-targeting or control shRNAs. Statistic assessment via two-tailed Student’s t test. **(N)** Visualization of PKM isoform expression in 134 patients of prostate tumors and matched normal tissues in CPGEA cohort by RNA-seq profiling [PMID: 32238934]. **(O and P)** Differential expression analysis of PKM1 (**O**) or PKM2 (**P**) in this patient cohort of 134 tumor-normal paired prostate specimens. The data was examined by the Wilcoxcon test compares two paired groups.

The Warburg effect, which altered metabolism by increasing the rate of glucose uptake and lactate production, provides a selective advantage for cancer tumorigenesis and progression. PKM2 isoform plays vital role in cancer cells by enhancing the Warburg effect ^33, 35, 36^. To test whether the ZMYND11 was also involved in the Warburg effect, we measured the extent of lactate production and glucose uptake in cells. The full-length ZMYND11 but not MYND-domain-deleted mutant significantly decreased the lactate production and glucose uptake **(Figures 5K and S5F)**, while silencing ZMYND11 resulted in the opposite observation **(Figures 5L and S5G)**. Furthermore, we analyzed the RNA splicing of the ZMYND11 knockdown prostate cell line 22Rv1. The PKM2 isoform was significantly increased after ZMYND11 knockdown **(Figure 5M)**. Increased PKM2 isoform was also observed in a cohort of 134 human prostate tumors compared to paired normal tissue samples ^32^ **(Figures 5N-5P)**. Taken together, these results showed that ZMYND11 suppressed PCa cell aerobic glycolysis by inhibiting HNRNPA1-dependent PKM splicing.

### ZMYND11 specifically binds the 194 arginine residues in the RGG motif of HNRNPA1

We next sought to gain molecular and biochemical insight into how ZMYND11 specifically binds to HNRNPA1. Structurally, HNRNPA1 contains two RNA recognition motifs (RRMs) and a glycine-rich Arg-Gly-Gly (RGG) domain at carboxyl (C)-terminus **(Figure 6A)**, possessing RNA-binding and protein-interaction capabilities, respectively ^7^. Our mass spectrometry analysis of the immunopurified HNRNPA1 revealed diverse posttranslational modifications, especially the frequent arginine methylation occurred in the RGG-motifs **(Figure S6A)**. Arginine methylation within RGG-motifs was known to mediate RNA binding ^37^, stress granule localization and formation ^23,38, 39^, yet whether and how this arginine methylation functioning in protein-protein interaction and recognition remains less defined. It is established that ZMYND11 is an epigenetic reader specifically recognizing H3.3K36me3 via its PWWP domain ^3,4^. Our results demonstrated that ZMYND11 is a dysregulated chromatin reader in cancers including PCa and could restrict HNRNPA1-mediated stress granule formation and oncogenic activity. Hence, we postulated that ZMYND11 may possess noncanonical function as a nonhistone reader and recognize methylated HNRNPA1 via its MYND reader module. We therefore first examined whether the RGG domain of HNRNPA1 interacts responsibly with ZMYND11, and accordingly generated HNRNPA1 mutation construct with RGG-domain deletion, co-expressing with V5-tagged ZMYND11 in HEK293T cells. This analysis showed that deletion of the RGG domain greatly diminished the binding of ZMYND11 to HNRNPA1 **(Figure 6B)**, indicating that the RGG domain of HNRNPA1 is essential for its interaction with ZMYND11. Given that six arginine residues in the RGG motif could be potentially methylated ^40^, we next examined which arginine methylation sites are involved in the interaction with ZMYND11. We therefore next generated single-site mutants by replacing arginine (R) with lysine (K), including R194K, R196K, R206K, R218K, R225K and R232K, and co-expressed them individually with ZMYND11. The results demonstrated that the R194K mutant greatly attenuated HNRNPA1 binding to ZMYND11 **(Figures 6C and S6B)**, indicating that the R194 residue of HNRNPA1 are paramount for the protein-protein interaction. Moreover, we assessed the potential functional role of R194K and found that the R194K mutant but not R206K consistently reduced colony formation, cell growth, migration, and invasion **(Figures 6D-6G and S6C),** despite that the R206K mutant also weakened the HNRNPA1-ZMYND11 interaction, supporting that the R194 residue is indispensable for tumor-promoting function of HNRNPA1. Taken together, these results indicated that ZMYND11 could specifically bind to the 194 arginine residue in the RGG motif of HNRNPA1.

**Figure 6:**
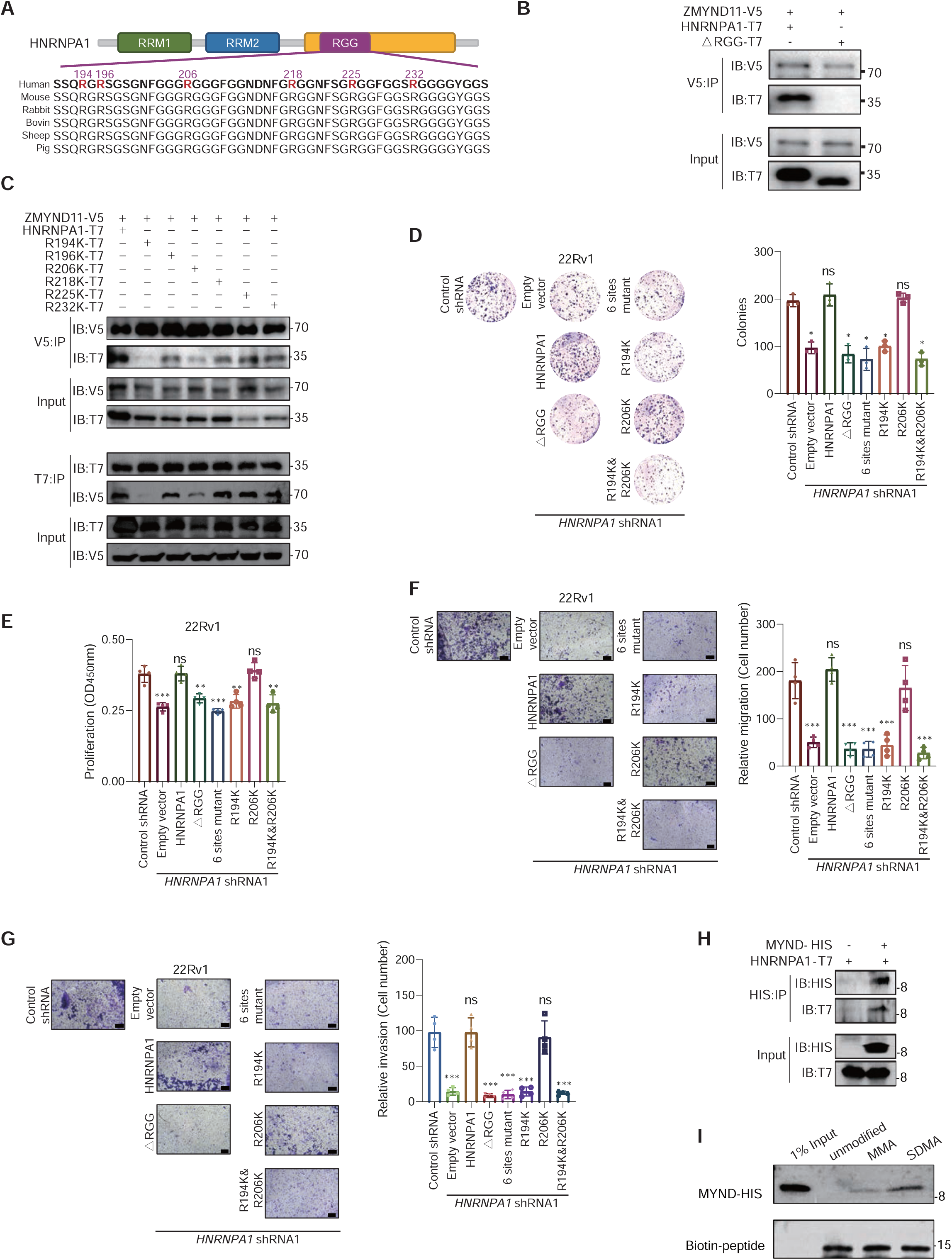
HNRNPA1 arginine 194 methylation is required for its oncogenic function and recognition by ZMYND11. **(A)** Schematic diagram showing the protein domain organization of HNRNPA1. Shown are amino acid sequence conservation of the glycine-arginine-rich (RGG) motif with highlighted arginine (R) residues (red) at the indicated position. RRM: RNA recognition motif. **(B)** Whole-cell lysates from HEK293T cells expressing V5-tagged ZMYND11 and T7-tagged HNRNPA1 or its mutant lacking the RGG-box were subjected to co-immunoprecipitation (co-IP) with anti-V5 antibody followed by immunoblotting (IB). **(C)** The lysates from HEK293T cells expressing V5-tagged ZMYND11 and T7-tagged HNRNPA1 or the indicated mutant constructs, were immunoprecipitated with anti-V5 antibody followed by immunoblotting (IB), and the reciprocal co-IP shown in the bottom. **(D-G)** Representative images and quantification of colony-formation (**D**), cell proliferation (**E**), migration (**F**), and invasion (**G**) of 22Rv1 cells with stable knockdown of HNRNPA1 and rescue by co-transfection with the indicated constructs. Scale bar, 200 μm. Statistical significance was assessed using two-tailed Student’s t test. ns: not significant, *: P<0.05, **: P<0.01, ***: P<0.001. **(H)** His-tag pull-down assay investigating direct interactions between the MYND domain of ZMYND11 and HNRNPA1. **(I)** Pull-down assays between the MYND domain of ZMYND11 and the biotin-labeled peptides carrying R194 monomethylation (MMA) or symmetric demethylation (SDMA).

### ZMYND11 specifically reads symmetric dimethyl arginine (SDMA) of HNRNPA1

Arginine methylation is a post-translational modification that is implicated in a variety of nuclear and cytosolic proteins, including histones, signaling molecules, and RNA splicing factors ^41^. The form of methylation of arginine residues can be classified into three: ω-NG-monomethyl-arginine (MMA), ω-NG, NG-asymmetric dimethyl arginine (ADMA), and ω-NG, NG-symmetric dimethyl arginine (SDMA) ^42^. To identify which arginine methylation in the R194 residues, we overexpressed T7-tagged wild-type HNRNPA1 and six arginine-mutated HNRNPA1 in 22Rv1 cells. Through immunoprecipitation (IP) of T7, we observed no difference in MMA, ADMA, or SDMA between wild-type HNRNPA1 with six single-arginine-mutated HNRNPA1 **(Figure S6D)**, suggesting that single residue mutation may have no effect to change the methylation. Thus, we constructed the mutants with all six arginine sites of HNRNPA1 and R194 unmutated HNRNPA1 for an independent IP assay. We observed that the R194 residue took the form of MMA and SDMA **(Figure S6E)**. To further examine the binding activity of ZMYND11 reader module directly to arginine methylated HNRNPA1, we purified recombinant protein for the MYND domain of ZMYND11. As shown in **Figure 6H**, the purified fragment could directly bind to ectopically expressed HNRNPA1 in a Co-IP experiment. Further pull-down assay confirmed the MYND domain of ZMYND11 specifically reading symmetric dimethyl arginine (SDMA) of HNRNPA1 in R194 site **(Fig.6I)**. Taken together, these results indicated that ZMYND11 could specifically recognize symmetric dimethyl arginine (SDMA) of HNRNPA1.

### ZMYND11 specifically reads SDMA of HNRNPA1 via the 572 tyrosine residue

To gain insight into the molecular basis underlying ZMYND11-MYND domain mediated recognition of HNRNPA1, we analyzed the crystal structure of ZMYND11-MYND and Epstein-Barr virus (EBV) nuclear antigen 2 (EBNA2) complex (PDB:5HDA) to predict specific amino acid residues of ZMYND11 in responsible for binding to HNRNPA1 ^43^. As a result, five residues in the MYND domain of ZMYND11: Y564, Y572, Y580, Q586 and W590, which are important for binding to EBNA2, could potentially contribute to HNRNPA1 recognition **(Figures 7A and 7B).** We then performed mutagenesis assays by generating single-site mutants while replacing them with alanine (A), and observed that mutation of Y572A in ZMYND11 greatly diminished its binding to HNRNPA1 **(Figures 7C and S7A)**, indicating that the ZMYND11 Y572 residue is paramount for its interaction with HNRNPA1. Consistently, our findings were further validated through two independent pull-down assays, confirming that the purified ZMYND11 Y572A mutant protein failed to recognize the SDMA modifications on HNRNPA1 **(Figures 7D and S7B)**. Furthermore, we observed that the Y572 mutant rescued the WT ZMYND11 inhibitory phenotype on colony formation, cell growth, migration, or invasion in PCa cells, suggesting that the Y572 residue may play a pivotal role in conferring the tumor-suppressive function of ZMYND11 **(Figures 7E-7H)**. As expected, WT ZMYND11, but not the Y572A mutant inhibited PKM2 and promoted PKM1 isoform formation both at protein **(Figure 7I)** and mRNA levels **(Figure 7J)**. The Y572A mutant had no effect on lactate production and glucose uptake **(Figures 7K and S7C)**. In addition, the immunofluorescence analysis revealed that the stress-induced cytoplasmic accumulation of HNRNPA1 was substantially attenuated by WT ZMYND11 but not the Y572A mutant in response to SA treatment **(Figure 7L).** Taken together, ZMYND11 could specifically read SDMA of HNRNPA1 via the Y572 residue, which was responsible for regulating PKM splicing and restricting cytoplasmic relocalization of HNRNPA1 under stress.

**Figure 7:**
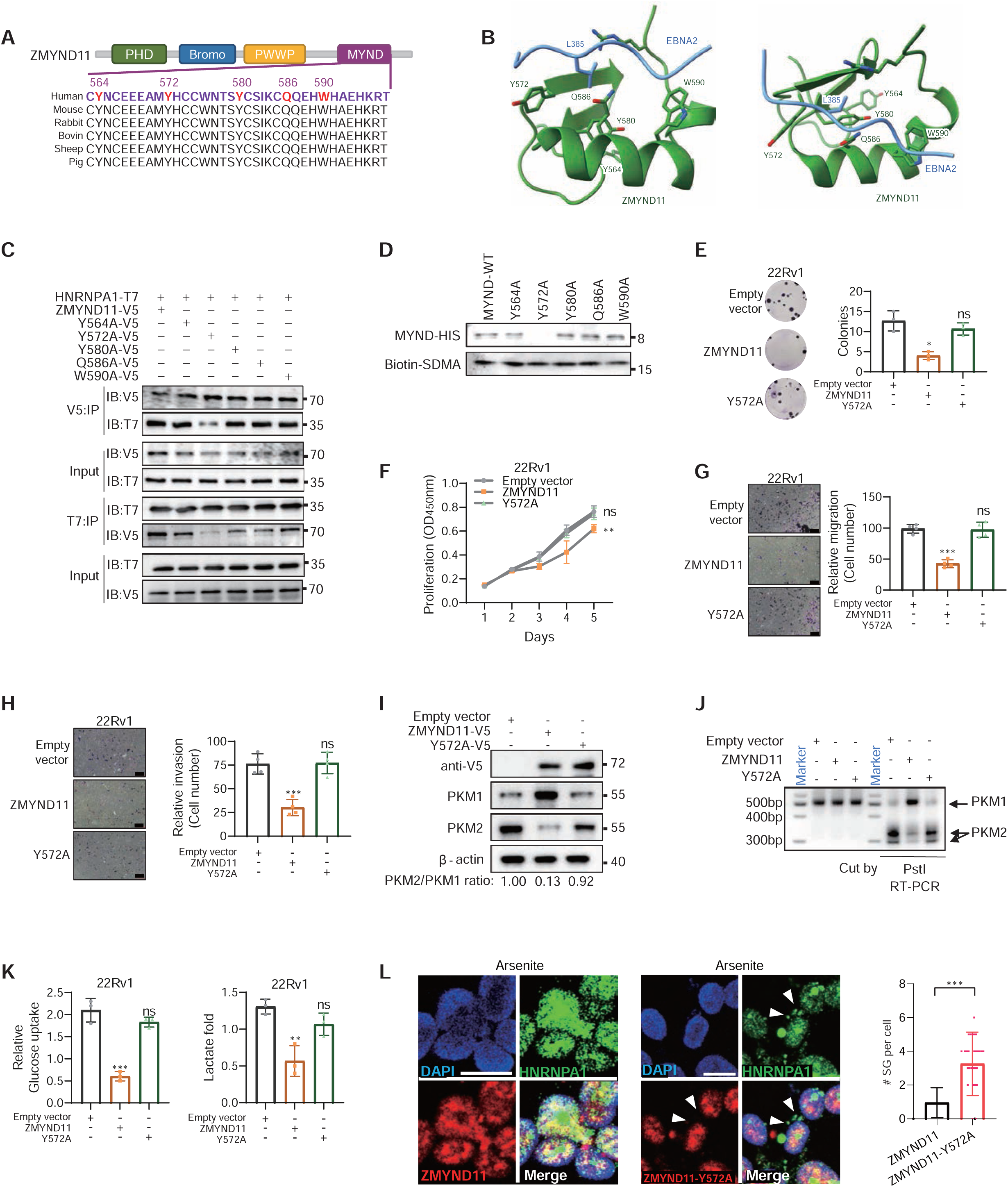
Tyrosine residue 572 of ZMYND11 is functionally important and essential for specifically reading SDMA on HNRNPA1. **(A)** Schematic representation of the MYND domain of ZMYND11. Shown is MYND domain organization along with potential candidate amino acid residues likely to be involved in binding with HNRNPA1. MYND domain conserved across species is shown. **(B)** Two views of the crystal structure (PDB:5HDA) for the MYND domain (green) of ZMYND11 complexes with EBNA2 (blue). Potential candidate amino acid residues responsible for recognition with HNRNPA1 are show in sticks. **(C)** Co-immunoprecipitation assay in HEK293T cells expressing T7-tagged HNRNPA1 and V5-tagged ZMYND11 or the indicated mutants. **(D)** Pull-down assays demonstrated protein-protein interactions between HIS-tagged MYND domain of ZMYND11 or the mutants and the biotinylated peptide containing R194 SDMA. **(E-H)** Representation and quantification of colony-forming (**E**), cell proliferation (**F**), migration (**G**), and invasion (**H**) of 22Rv1 cells overexpressing ZMYND11 or the Y572A mutant. Scale bar, 200 μm. *P<0.05, **P<0.01, ***P<0.001, were examined by two-tailed Student’s t test. **(I-J)** PKM splicing assay (**I**) and immunoblots for the indicated PKM isoforms **(J)** were performed in 22Rv1 cells ectopically expressing ZMYND11 or the Y572A mutant. **(K)** Relative glucose uptake and lactate production were measured in 22Rv1 cells with an ectopical expression of ZMYND11 or Y572A mutant. ns, not significant, *P<0.05, **P<0.01, ***P<0.001, two-tailed Student’s t test. **(L)** Fluorescence colocalization microscopy analysis of ZMYND11 or the Y572A mutant with HNRNPA1 in 22Rv1 cells treated with sodium arsenite or control. Note that stress-induced cytoplasmic accumulation of HNRNPA1 was substantially attenuated by wild-type ZMYND11, but not the Y572A mutant. Arrows, stress granules in cells. Mean ± SD, was assessed using two-tailed Student’s t-test. *** represents p < 0.005. Scale bars, 25 μm.

### Depletion of PRMT5 impairs the interaction of ZMYND11 and HNRNPA1

The methylation of arginine residues is catalyzed by the family of arginine methyltransferases (PRMTs), which are classified into type I, II, and III through the methyl-arginine products. Type I, including PRMT1, 2, 3, 4, 6, 8, catalyze monomethylarginine (MMA) and asymmetric dimethylarginine (ADMA) ^44^. PRMT5 and 9 belong to type II, and catalyze MMA and symmetric dimethylarginine (SDMA). PRMT7, a type III PRMT, catalyzes only MMA **(Figure 8A)**. Our results indicated that the methylation of HNRNPA1 R194 was mainly SDMA **(Figure 6I)**. To further elucidate which PRMTs regulate the methylation of HNRNPA1, we performed a Co-IP assay by co-expressing T7-tagged HNRNPA1 and nine GFP-PRMTs in HEK293T cells, leading to the finding of PRMT1 and PRMT5 with higher binding affinity towards HNRNPA1 **(Figure 8B)**. In line with the data, PRMT1, PRMT5 and PRMT7 were reported to regulate arginine methylation of HNRNPA1 ^40^. We therefore generated individual PRMT1/5/7 knockdown cell models to test the effect of PRMT-mediated HNRNPA1 arginine methylation on the association with ZMYND11 **(Figure S8A)**. The results showed that knocking down PRMT5 greatly compromised the interaction between HNRNPA1 and ZMYND11, whereas their interaction was not influenced in the cells having PRMT1 or PRMT7 knockdown **(Figures 8C and S8B)**. Furthermore, we observed a disruption of the interaction between endogenously expressed ZMYND11 and HNRNPA1 in 22Rv1 or DU145 cells with stable knockdown of PRMT5 **(Figures 8d and S8C)**. Consistent with the observations, the protein-protein interaction of HNRNPA1 and ZMYND11 was greatly diminished when treated the cells with PRMT5 inhibitors (EPZ015666 or GSK3326595) while the inhibitory effects were substantially weaker for the PRMT1 inhibitors AM1 and DCLX069 **(Figures 8E and 8F)**. Taken together, our results demonstrated that ZMYND11 specifically read SDMA of HNRNPA1 while PRMT5 was responsible for the arginine methylation and essential for the interaction between ZMYND11 and HNRNPA1.

**Figure 8:**
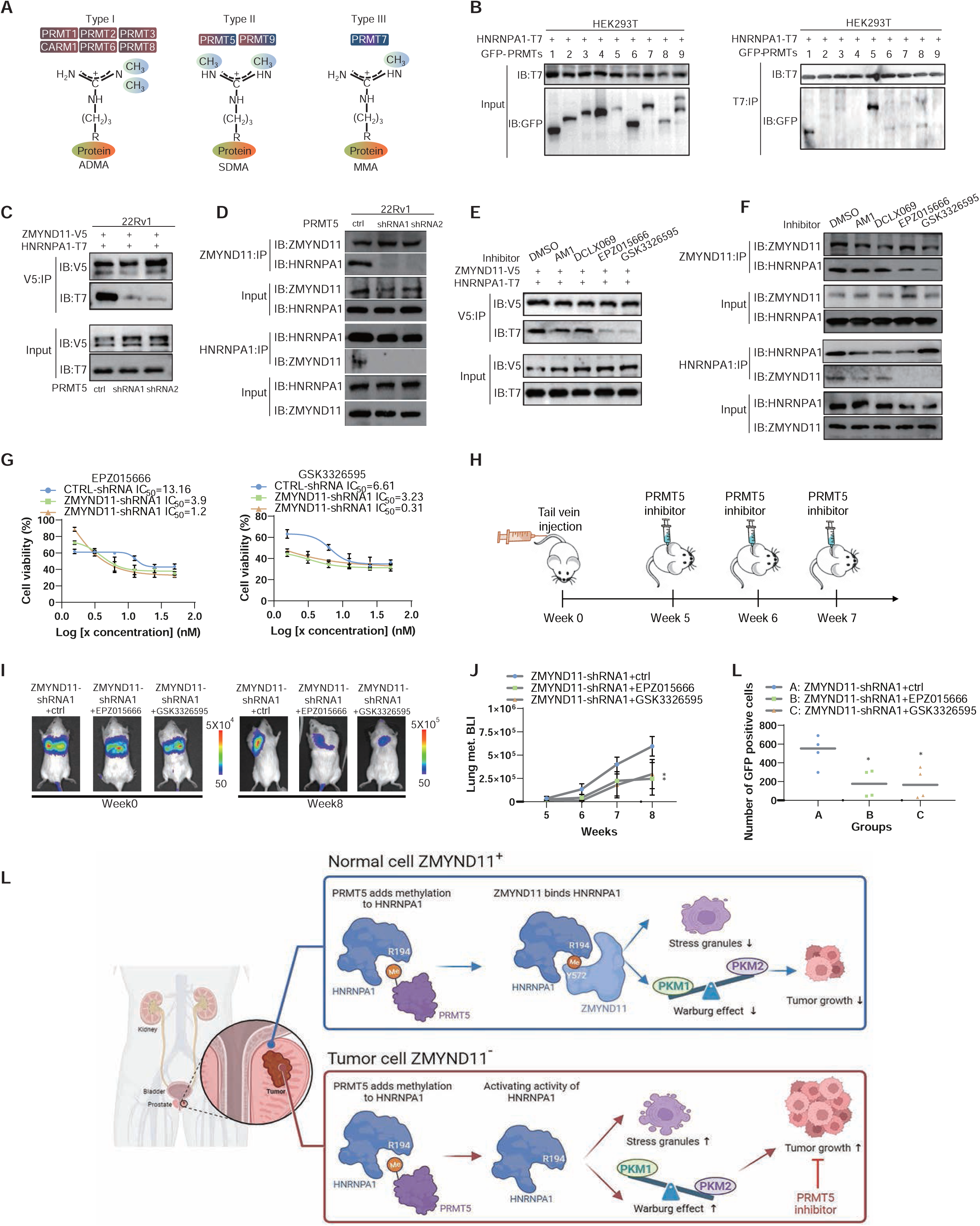
Pharmacological inhibition of PRMT5 impairs ZMYND11-HNRNPA1 interaction and suppresses in vivo metastatic capacity of PCa cells with ZMYND11-low expression. **(A)** Schematics of arginine methylation states by PRMT family of enzymes. **(B)** Co-immunoprecipitation assay of HEK293T cells expressing T7-tagged HNRNPA1 and the indicated GFP-tagged PRMT family member. **(C)** Co-immunoprecipitation assays using 22Rv1 cell protein extracts co-expressing V5-tagged ZMYND11 and T7-tagged HNRNPA1 while having stable expression of control or shRNAs against PRMT5. **(D)** Immunoprecipitation-Western blot analysis of an endogenous interaction between ZMYND11 and HNRNPA1 in 22Rv1 cells stably expressing control or ZMYND11-targeting shRNAs. **(E)** Immunoprecipitation using V5-tag antibodies from 22Rv1 cells ectopically co-expressing ZMYND11 and HNRNPA1 while treated with PRMT1 inhibitors (AM1 or DCLX069, 50μM, 48h) or PRMT5 inhibitors (EPZ015666 or GSK3326595, 10μM, 48h). **(F)** Co-immunoprecipitation assay of an endogenous interaction between ZMYND11 and HNRNPA1 in 22Rv1 cells treated with the indicated PRMT1/5 inhibitor. **(G)** Half-inhibitory concentration (IC50) test of two PRMT5 inhibitors (EPZ015666 or GSK3326595, 10μM, 48h) for ZMYND11 knockdown or control cells in 22Rv1. **(H)** Schematics of *in vivo* mouse experiments with PRMT5-Selective Inhibitors. **(I and J)** 22Rv1 cells with stable knockdown of ZMYND11 while expressing luciferase were injected into the tail-vein of male NOD-SCID mice. At 5, 6 and 7 weeks after injection, mice were subject to intraperitoneal treatment with the indicated PRMT5 inhibitors. Shown are the representative images at indicated time points (**I**) and weekly quantification of BLI photon flux of lung metastasis in mice (**J**). Errors bar, ±SD, n = 5. *p < 0.05, **p < 0.01, ***p < 0.001, Student’s t tests. **(K)** FACS-analysis of GFP-positive 22Rv1 cells (CTCs) in the peripheral blood of SCID mice (n=4). The scatter plot indicated the number of CTCs recovered from each mouse treated with vesicles or PRMT5 inhibitors. Statistical significance was assessed using two-tailed Student’s t test. * p < 0.05. **(L)** A model for ZMYND11 recognition of arginine methylation restrains HNRNPA1-mediated tumor progression. Top: higher expression of ZMYND11 in normal cells could specifically recognize R194 methylation of HNRNPA1 catalyzed by PRMT5, thereby compromising the role of HNRNPA1 in stress granule formation and PKM splicing for tumor aggressiveness. Bottom: in tumors with lower expression of ZMYND11, HNRNPA1 ensures an increased formation of stress granule and a higher PKM2/PKM1 ratio contributing to tumor progression. PRMT5 inhibitors are promising in attenuating cancer progression.

### Pharmacological inhibition of PRMT5 suppresses the growth of ZMYND11 low-expressing tumor

Given the critical role and clinical relevance of PRMT5 inhibitors in cancer progression ^45, 46^, we sought to evaluate whether PRMT5 inhibitors could be an effective strategy for the treatment of ZMYND11 low expression PCa. As expected, PRMT5 inhibitors (EPZ015666 or GSK3326595) greatly suppressed cell growth, colony formation, migration and invasion in ZMYND11 knockdown cells. However, inhibitors did not have a significant effect in control cells **(Figures S8D-S8F)**. Half-inhibitory concentration (IC50) test also confirmed that the inhibitors were more effective in killing the ZMYND11 knockdown cells than control **(Figure 8G)**. Remarkably, using PRMT5 inhibitors EPZ015666 or GSK3326595, we observed that lung metastasis was markedly suppressed in the bioluminescent imaging (BLI)-based lung metastasis mouse model after tail vein injection of the PCa cells with stable knock-down of ZMYND11 **(Figures 8H-8J)**. GFP-positive circulating cancer cells in the blood were obviously reduced upon the treatment with inhibitors **(Figure 8K)**. These data indicated that the pharmacological inhibition of PRMT5 was effective in suppressing the growth of PCa cells and tumors, and ZMYND11 might serve as a predictive biomarker to optimize the clinical management of patients.

## Discussion

It has been well studied that HNRNPA1 plays a pivotal role in upregulating the PKM2/PKM1 ratio through alternative splicing, resulting in subsequent enhancement of aerobic glycolysis, while increased expression of HNRNPA1 was driven by the well-established oncogenic transcription factor c-Myc ^8^. Despite substantial overexpression of MYC, HNRNPA1, and PKM2 in most tumors ^8, 47^, it is noteworthy that the MYC-HNRNPA1-PKM2 axis is not universally responsible for driving tumorigenesis and tumor progression in all cancer types. This intriguing observation hints at the existence of inhibitory factors that restraint the functionality of this regulatory axis. In this study, we revealed that the epigenetic reader ZMYND11 possesses a newly-identified noncanonical function as nonhistone methyl reader and could recognize arginine methylation of HNRNPA1 at arginine 194, and thereby downregulates HNRNPA1-mediated PKM splicing and aerobic glycolysis **(Figure 8L)**. Our study highlights the potential of ZMYND11 to restrict the oncogenic activity of HNRNPA1-mediated PKM splicing and stress granule formation in cancer cells. Consistently, we provide clinical evidence in supporting a reverse association between ZMYND11 expression and tumor progression to metastasis, higher tumor stages as well as a poorer clinical prognosis. These results underscore the potential significance of ZMYND11 as a valuable biomarker for aiding clinical diagnosis and treatment in certain cancer context.

ZMYND11 was previously identified as an epigenetic reader, specifically binding to histone H3.3K36me3, and modulating RNA Pol II during the elongation stage and thus gene expression suppression ^2, 3^. Our study revealed a novel role of ZMYND11, showing its capacity of independently regulating HNRNPA1 and its downstream networks while having PRMT5-induced arginine methylation. This noncanonical function of ZMYND11 furthers our understanding of its tumor-suppressive mechanisms and intricate crosstalk with the oncogenic regulatory axis such as HNRNPA1-PKM2. Furthermore, we propose a model in which ZMYND11, led by its MYND domain rather than the more conventional ‘epigenetic reader’ modules PBP, recognizes the methyl-modified nonhistone protein HNRNPA1. To be more precise, it is the Y572 site within the MYND domain that plays this pivotal role. Collectively, our findings provide novel insights into the tumor-suppressor mechanism of ZMYND11, attributed to its previously unknown nonhistone methylation reader function **(Figure 8L)**.

HNRNPA1 proteins can form stress granules (SGs) under stress, which are dynamic non-membrane compartments assembled temporarily in response to cellular stress, and predict a poor prognosis ^10, 15, 16^. Asymmetrically and symmetrically dimethylated RGG domains are commonly known to induce phase separation ^48, 49^. The mechanisms of SG formation remain unclear. Here, we identified the MYND domain of ZMYND11 specifically recognized symmetrically dimethylated RGG motif of HNRNPA1, limiting HNRNPA1 cytoplasmic translocation and thus dampening the formation of SGs under stress **(Figure 8L)**. Studies have reported that multiple chemotherapeutic drugs can trigger the assembly of SGs, potentially contributing to chemotherapy resistance, including 5-Fluorouracil, Arsenic trioxide (ATO), paclitaxel, and others ^50–52^, which suggests that targeting HNRNPA1 or ZMYND11 might overcome cancer resistance.

We reported that the SDMA modifications on HNRNPA1 RGG domains are catalyzed by PRMT5. PRMT5 inhibitors are being pursued as a promising cancer treatment approach, and numerous mechanisms have been proposed for the efficacy of PRMT inhibition ^45, 46^. Our observations demonstrated that PRMT5 inhibitors effectively repressed ZMYND11-low-expressing cells *in vitro* and tumor progression *in vivo*. In this scenario, ZMYND11 could serve as an excellent marker to guide clinical diagnostics and therapy in cancer, especially when considering the use of PRMT5 inhibitors. Notably, based on the ongoing clinical trials with PRMT5 inhibitor (NCT03614728), our findings may potentiate its therapeutic implications for the cancer patients with lower expression levels of ZMYND11. Follow-up work warrants more clinical samples before and after PRMT5 inhibitor treatment to test rigorously this hypothesis in proper cohorts of cancer patients.

## Methods

### Mouse experiments

Animal studies mentioned in the context were conducted in compliance with the guidelines for the care and use of laboratory animals and were approved by the Institutional Animal Care and Use Committee of Fudan university. This study describes the subcutaneous tumor model in which 1×10^7^ 22Rv1 cells in 100μL of PBS were injected into the left flanks of 5-week-old male BALB/c nude mice. After six weeks, the mice were euthanized, and the tumor volumes were calculated. For the spontaneous lung metastasis model, 2×10^6^ 22Rv1 cells in 100 μL of PBS were injected into the tail vein of 5-week-old male NOD-SCID mice. After 6 weeks, these mice were sacrificed and examined for tumor growth and metastasis in the lung using an *in vivo* imaging system (IVIS). The mice were randomly grouped based on similar body weights, and no mice were excluded from the analysis except for those that died unexpectedly due to reasons unrelated to the tumor. The investigators conducting the experiment were aware of the group allocation and assessed the outcomes without being blinded.

### Cell Lines

22Rv1, LNCaP, PC3, A549 were grown in RPMI1640 (Invitrogen) and DU145, MDA-MB-231 and HEK293T were grown in DMEM (Invitrogen). Cells were maintained at a temperature of 37°C in a 5% CO2 incubator. The base medium was supplemented with 10% fetal bovine serum (FBS, Giboc) and 1% of a combination of penicillin and streptomycin (BI) to provide additional nutrients and prevent bacterial contamination.

### Human prostate cancer tissues and non-cancer tissues

Paraffin-embedded prostate cancer tissues and non-cancer tissues were obtained from Fudan University Shanghai Cancer Center with informed consent from all subjects and approval from the Hospital Institutional Research Ethics Committee.

### Quantitative RT-PCR

The RNA was isolated from cultured cell lines or mouse tissues using the RNeasy Mini Kit from QIAGEN. This kit allows for efficient isolation of high-quality RNA from various sources.

To synthesize cDNA from the isolated RNA, the High-Capacity cDNA Reverse Transcription Kit from TAKARA was used. This kit includes all the necessary reagents and enzymes for efficient reverse transcription of RNA into cDNA. For quantitative RT-PCR analysis, the SYBR Select Master Mix from TAKARA was used. This master mix contains all the components required for quantitative PCR, including the SYBR Green dye that allows for real-time monitoring of amplification. Primer sequences used in this experiment can be found in Table.

### Western blot

The denatured samples were loaded onto a polyacrylamide gel and separated by electrophoresis. The proteins were then transferred onto a nitrocellulose membrane and blocked with 5% skimmed milk. The membrane was incubated with primary antibody overnight at 4°C, followed by incubation with secondary antibody conjugated with a fluorescent or enzymatic tag. The protein bands were visualized using a chemiluminescent or fluorescent detection system. The intensity of the bands was quantified using image analysis software. The nuclear and cytoplasmic separation experiments were performed with the kit of NE-PER (Thermo Fisher Scientific, 78833).

Antibodies used are as follows: Anti-ZMYND11 antibody (CST, 677135); Anti-HNRNPA1 (proteintech, 67844-1-Ig); Anti-V5 tag monoclonal antibody (Thermo Fisher Scientific, R96025), HRP-anti-V5 Tag monoclonal antibody (Thermo Fisher Scientific, R96125); Anti-human GAPDH (Proteintech, 10494-1-AP); HRP-Goat anti-mouse IgG (Proteintech, SA00001-1); HRP-Goat anti-Rabbit IgG (Proteintech, SA00001-2); Anti-Beta Actin antibody (Proteintech, 66009-1-Ig); Anti-PKM1 Polyclonal antibody (Proteintech, 15821-1-AP); Anti-PKM2 polyclonal antibody (Proteintech, 15822-1-AP); Anti-T7 tag monoclonal antibody (Bethyl, A190-117A); Anti-PRMT1 monoclonal antibody (Santa cruz, sc-166963); Anti-PRMT5 monoclonal antibody (Santa cruz, sc-376937); Anti-PRMT7 rabbit polyclonal antibody (Abclonal, A12159); Anti-G3BP1 rabbit polyclonal antibody (Abclonal, A14836); Anti-His-Tag monoclonal antibody (Abclonal, AE003).

### siRNA transfection

50%–60% confluent 22Rv1 cells or 60%–70% confluent LNCaP cells were seeded in 6-well plates. 24 h later, lipofectamine RNAiMAX reagent (Invitrogen) was diluted by Opti-MEM, and siRNAs were also diluted by Opti-MEM. The diluted siRNAs were added into diluted lipofectamine RNAiMAX reagent, incubating at room temperature for 10 min. Then, the siRNA-lipid complexes were added to cultured cells. Medium was changed after 24 h and the cells were collected after 48 h.

### Lentiviral constructs, lentivirus production and infection

The shRNA constructs were ordered from Functional Genomics Unit (University of Helsinki) or designed by BLOCK-iT RNAi Designer (Thermo Fisher Scientific) and inserted into pLKO.1 Vector (Addgene). To package the third generation lentiviral vectors, 293T cells were used. In detail, 293T cells were grown to 65%-75% confluency in 6 cm dishes. The cells were trypsinized and seeded into new 6 cm dishes. After 24 hours, the growth medium was replaced with fresh FBS-free DMEM (Invitrogen) supplemented with 0.1% penicillin and streptomycin. The cells were co-transfected with the following plasmids:

- Indicated shRNA construct or overexpression construct (3 μg each)
- pVSVG (envelope plasmid, 1 μg)
- pMDLg/pRRE (packaging plasmid, 1 μg)
- pRSV-Rev (packaging plasmid, 1 μg)

The transfection was performed using 18 μL of PEI (polyethyleneimine). After transfection, the medium was replaced with fresh medium. Every 12 hours, the virus-containing medium was collected up to six times. The collected lentivirus was passed through a 0.45 mm filter unit to remove cellular debris. The filtered lentivirus was snap frozen in liquid nitrogen. The frozen lentivirus was stored at −80°C for future use.

The lentivirus-mediated knockdown was performed by infecting the target cells with lentivirus containing the necessary shRNA sequence. After 24 hours, the virus was removed and replaced with normal growth medium containing 1 mg/ml puromycin from Sigma. The puromycin selection was used to kill off any uninfected control cells. Once the uninfected control cells had completely died, the target cells that were successfully infected with the lentivirus were cultured in normal growth medium supplemented with a lower concentration of puromycin, specifically 0.5 mg/ml. This lower concentration was sufficient to maintain the selection pressure on the target cells.

For lentivirus-mediated overexpression, 22Rv1, LNCaP and DU145 cell lines were generated by lentiviral transduction, cloned into pcDNA3.1 (Addgene) and pCDH (Addgene).

### Co-IP and mass-spectometry (MS) analysis

Cultured cells were rinsed with pre-cooled PBS and lysed by IP buffer (glycerophosphate, 1.5 mM MgCl_2_, 2 mM ethylenebis (oxyethylenenitrilo) tetraacetic acid (EGTA)). Cell lysates were centrifuged at 10,000 × g for 15 min at 4°C to remove intact cells. The supernatant was either incubated with control immunoglobulin G (IgG) or primary antibody overnight in IP buffer with protease inhibitors), followed by incubation with 20 μL of resuspended volume of Protein G beads (Roche) for 2 h at 4 °C to pull down bound proteins. Beads were centrifuged at 1000 × g for 5 min at 4°C to remove the supernatant, washed four times with the IP buffer and boiled for 10 min at 95°C. Samples were run on SDS PAGE gel, followed by Coomassie Brilliant Blue R-250 (Bio-Rad) staining. Afterwards, gel bands were excised, destained, trypsinized and subjected to MS analysis to identify individual proteins using liquid chromatography-Tandem Mass Spectrometry LC-MS/MS (Orbitrap Velos Pro mass spectrometer, Thermo Fisher Scientific). The mass spectrometry analysis of posttranslational modification uses the same method and instrument.

### RNA-seq library generation and sequencing

Total RNA of each sample was extracted using TRIzol Reagent (Invitrogen)/RNeasy Mini Kit (Qiagen)/other kits. Total RNA of each sample was quantified and qualified by Agilent 2100 Bioanalyzer (Agilent Technologies, Palo Alto, CA, USA), NanoDrop (Thermo Fisher Scientific Inc.) and 1% agrose gel. 1 μg total RNA with RIN value above 6.5 was used for following library preparation. Next generation sequencing library preparations were constructed according to the manufacturer’s protocol of VAHTS mRNA-seq V3 Library Prep Kit for Illumina (NR611). The poly(A) mRNA isolation was performed using Poly(A) mRNA Magnetic Isolation Module or rRNA removal Kit. The mRNA fragmentation and priming were performed using First Strand Synthesis Reaction Buffer and Random Primers. First strand cDNA was synthesized using ProtoScript II Reverse Transcriptase and the second-strand cDNA was synthesized using Second Strand Synthesis Enzyme Mix. The purified double-stranded cDNA by beads was then treated with End Prep Enzyme Mix to repair both ends and add a dA-tailing in one reaction, followed by a T-A ligation to add adaptors to both ends. Size selection of Adaptor ligated DNA was then performed using beads, and fragments of ∼420 bp (with the approximate insert size of 300 bp) were recovered. Each sample was then amplified by PCR for 13 cycles using P5 and P7 primers, with both primers carrying sequences which can anneal with flow cell to perform bridge PCR and P7 primer carrying a six-base index allowing for multiplexing. The PCR products were cleaned up using beads, validated using an Qsep100 (Bioptic, Taiwan, China), and quantified by Qubit3.0 Fluorometer (Invitrogen, Carlsbad, CA, USA).

Then libraries with different indices were multiplexed and loaded on an NovaSeq6000 instrument according to manufacturer’s instructions (Illumina, San Diego, CA, USA). Sequencing was carried out using a 2×150 bp paired-end (PE) configuration; image analysis and base calling were conducted by the NovaSeq6000 Control Software (HCS) + OLB + GAPipeline-1.6 (Illumina) on the NovaSeq6000 instrument.

### RNA-seq data processing and bioinformatics analysis

RNA-seq data quality was assessed using FastQC (version: 0.11.9) (www.bioinformatics.babraham.ac.uk/projects/fastqc/). Low-quality reads were trimmed and adapters were removed using AdapterRemoval (version: 2.3.2) ^53^. Clean data were mapped to the reference genome (hg38) using STAR (version: 2.7.9a) ^54^ and the aligned BAM files were sorted using SAMtools (version: 1.13) ^55^. Next, our analysis followed the LeafCutter ^56^ pipeline in order to quantify the alternative splicing (AS) events. The interested differential splicing events were visualized using LeafViz application in LeafCutter ^56^.

### Cell viability and proliferation assays

The resuspended cells were carefully transferred into each well of a 96-well cell culture plate (4X 10^3^ for 22Rv1, 2X 10^3^ for LNCaP, 1X 10^3^ for DU145 per well, respectively). Cell viability and proliferation were determined using Cell Proliferation Kit II. The data were collected at specific time points by measuring the absorbance at 450 nm, following the instructions provided by the manufacturer. The values obtained from triplicate wells were used to calculate statistical significance using a two-tailed Student’s t-test.

### Colony formation assays

The resuspended cells after trypsinization were cultured in 6-well plates (1X10^3^ for 22Rv1 and LNCaP, 200 for DU145 per well, respectively), and colonies were allowed to develop over 7 to14 days. Following fixation with 3.7% paraformaldehyde and staining with crystal violet, visible colonies were quantified. Colonies containing at least 50 cells were counted manually or analyzed with image software. Results were expressed as mean colonies ± standard deviation. Experiments were performed in triplicate and statistical significance was determined using two tailed Student’s t-test.

### Invasion and migration assays

The cells were detached from the culture dish using trypsin and suspended in growth medium without serum or growth factors. The cell concentration was adjusted to 4X10^5^ cells/ml 22Rv1 or LNCaP, 2X10^5^ cells/ml DU145. 200 μL of the cell suspension was then transferred into 8-mm Transwell inserts, with or without a 100 μL coating of Matrigel. The Matrigel was diluted with serum-free medium to a concentration of 250 μg/ml. The lower chambers of the Transwell inserts were filled with 700 μL of normal growth medium. After a 36-hour incubation period, the cells were fixed using 3.7% formaldehyde and permeabilized with methanol. They were then stained using the Wright-Giemsa stain. Cells on the upper surface of the membranes were removed using a cotton swab. The invasive cells that migrated to the bottom surface of the filters were quantified by counting the number of cells that penetrated the membrane in twelve microscopic fields per membrane. The counting was done at 20X magnification. Statistical analysis was performed using a two-tailed Student’s t-test, comparing the results from three replicate inserts.

### PKM splicing assays

PKM splicing was performed as previously described (David et al., 2010) ^8^. In this protocol, total RNA was extracted using TRIzol. Reverse transcription of RNA was carried out using the PrimeScript RT reagent Kit with gDNAEraser from TAKARA. The resulting cDNA from reverse transcription was then amplified by PCR and digested using PstI. Primer sequences were PKM-F: 5’-CTGAAGGCAGTGATGTGGCC-3’; PKM-R: 5’-ACCCGGAGGTCCACGTCCTC-3’Finally, the digested mixtures were separated by 8% nondenaturing polyacrylamide gel electrophoresis (PAGE).

### Measurement of lactate production

Cells were transfected with the indicated siRNAs or plasmids for 36 h and then incubated in phenol red-free RPMI1640 medium without FBS for 4 h. The accumulation of lactate in the media was detected using a Lactate Colorimetric Assay kit II (K627-100, BioVision, USA). The background was corrected by deducting the OD value from the fresh phenol red-free medium RPMI1640. According to the lactate standard measurement values, the standard curve of nmol/well on OD450 nm was drawn. The sample OD450 nm values were applied to the standard curve to calculate the concentrations of lactate in test samples.

### Glucose uptake assay

Cells were transfected with the indicated siRNAs or plasmids for 36 h and incubated in DMEM medium without L-glucose and phenol red for 8 h. The amount of glucose in these media was measured using a Glucose Colorimetric Assay kit (K606-100, Bio Vision, USA). Fresh DMEM medium was used for the negative control. Three biological replicates were performed.

### Recombinant protein purification

BL21 (DE3) competent *E. coli* cells were used to express His-tagged recombinant proteins with the pET-28a plasmids. The single colony was cultured in LB liquid medium with 100 μg/ml kanamycin in 1L LB. The cells were shaken at 200 rpm 37 °C. IPTG with a final concentration of 1mM was added after reaching the exponential growth period of OD600 nm at 0.6 and incubated at 16 °C for 14 hours. To harvest the cells, the pellet was centrifuged at 10000 g for 10 minutes. The cells were treated with lysis buffer (0.3 M NaCl, 50 mM NaH_2_PO_4_ pH 8.0) with 1 mg/mL Lysozyme and sonicated for 5 minutes. Then the cells were centrifuged at 10000 g for 30 minutes and the supernatant was collected and filtered with a 0.45 μm filter. The protocol of purification was used with the HisSep Ni-NTA 6FF Chromatography Column (20504, Yeasen Biotech) following the manufacturer’s instruction. The target protein was dialyzed and washed with imidazole in PBS. Proteins were stored in a −80 °C refrigerator.

### Pull-down assay

Biotin-labeled peptides carrying R194 monomethylation (MMA); symmetric demethylation (SDMA) or unmodified fragment (Nanjing peptide biotechnic) was incubated with HIS-MYND fusion protein respectively at 4°C overnight. After incubating the mixture with Dynabeads (Thermo Fisher Scientific) for another 2 h at 4°C, beads were separated by magnetic frame and washed four times with 0.3% NP-40 buffer in PBS. Samples were boiled in SDS loading buffer and subjected to western blotting analysis.

### Half maximal inhibitory concentration (IC50) assay

IC50 values were obtained at several pre-incubation time points for two inhibitors of PRMT5, EPZ015666 (MCE) and GSK3326595 (MCE). 22Rv1 cells were seeded at 7000 cells per well of a 96-well cell culture plate and incubated for 24 h to allow for attachment. Cells were treated with PRMT5 inhibitors (EPZ015666 and GSK3326595, respectively) and DMSO as control of a concentration gradient (1.5625 nM, 3.125 nM, 6.25 nM, 12.5 nM, 25 nM, 50 nM) for 72 h. Following incubation with drugs, assay plates were removed from media, added the CCK-8 agent, and incubated at 37°C for 4 h. The plates were measured on a spectrophotometer at 450 nm.

### Immunohistochemistry (IHC)

In this study, immunohistochemistry (IHC) was performed using a standard LSAB protocol from Dako, Carpinteria, CA. The sections were stained for ZMYND11 using an anti-ZMYND11 rabbit monoclonal antibody (1:100, Abcam, USA), for HNRNPA1 using an anti-HNRNPA1 rabbit monoclonal antibody (1:100, proteintech, China).

### Immunofluorescence staining and stress granule analysis

Cells were seeded on coverslips in 24-well plates and were fixed with 4% paraformaldehyde for 15□min when growing to 60% confluence. 0.3% Triton X-100 containing PBS was used to permeate the cell membrane at room temperature for 10□min. The cover slides were then blocked in 3% BSA/PBS at room temperature for 1□h, and incubated with the primary antibody and fluorescent-labeled secondary antibody overnight at 4□°C or for 1□h at room temperature, respectively. Next, the nucleus was stained by DAPI in mounting buffer (Beyotime Biotechnology, P0131). Observation and photographing were performed with the Leica TCS SP8 confocal microscope, and image processing and analysis were performed with Leica TCS SP8 edition software.

Cells were treated with sodium arsenite (SA) (0.5mM, 1h, Sigma) to induce the formation of stress granules. Stress granules were manually quantified using the indicated markers G3BP1 (Abclonal, A14836, 1:100). At least 50-100 cells were counted for each condition in each experiment. Each group was taken three images and were then counted.

### Clinical analysis

The score of protein expression of ZMYND11 and HNRNPA1 of Fudan cohort was conducted depending on the staining area of IHC: 0, 0-5%; 1, 5-25%; 2, 26-50%; 3, 51-75%; and 4, > 75%, and the staining intensity was categorized as follows: no staining scored 0, weakly staining scored 1, moderately staining scored 2 and strongly staining scored 3, respectively. Composite expression score ^57^ is calculated from intensity and area measurements for immunostaining (CES = 4*(intensity score - 1) + area score), yielding a series of results ranging from 0 to 12. Tongji cohort samples were scored with the scoring software PE Vectra3 provided by the Olympus VS120 system, with a maximum score of 300 points. Alternative splicing assay of Changhai cohort of 134 prostate cancer tissues and matched adjacent NT tissues was used LeafCutter software.

### Survival analysis

Kaplan-Meier survival analysis was applied to assess the impact of ZMYND11 or HNRNPA1 on PCa prognosis in multiple independent cohorts. Patients were first stratified into two groups based on the median gene expression levels. For the investigation of the synergistic effect of ZMYND11 and HNRNPA1 on PCa patient survival, we first stratified patients based on the median expression level of ZMYND11 or HNRNPA1 and then selected patients with simultaneous high ZMYND11 & low HNRNPA1 or low ZMYND11 & high HNRNPA1 expression levels. Kaplan-Meier survival analysis was conducted using R package “Survival” (v. 3.2.3) and assessed by using log-rank test.

### Statistical analyses

The statistical analyses and data presentation in this study were conducted using GraphPad Prism 8.0 software from GraphPad Software. The specific details of the analyses and presentation can be found in the figure legends. In this study, statistical significance was determined by P values less than 0.05. The experiments conducted in vitro were repeated multiple times independently, and the results were consistently similar, as indicated in the figure legends. Statistical tests for investigating gene expression across normal, tumor, and metastatic tissues or clinical features, including Gealson score, tumor stage, or PSA were evaluated by the Mann-Whitney U test or the Kruskal-Wallis H test according to the number of comparison groups. For the results from microarray-based expression profiling, gene probes with lowest P values were selected. Samples with missing gene expression or patient survival data were excluded from analyses. Statistical tests were performed using RStudio (v. 1.4.1106) with R version v. 4.1.0.

## Data availability

We have used the following public datasets in this study: GSE21019, GSE6919, GSE3325, GSE6099, GSE21034, and TCGA PRAD. In addition, data sets used for investigating the ZMYND11 expression level in pan-cancer in **Figure 1a** include GSE7696 (Brain), GSE7803 (Cervix), GSE20842 (Colorectal), GSE23400 (esophagus), GSE13911 (Gastric), GSE2549 (Mesothelioma), GSE15471 (Pancreas), and GSE21034 (Prostate). The CPGEA RNA-seq data were acquired from http://www.cpgea.com/. All other types of data in supporting the discoveries in this study are available upon reasonable request for the corresponding authors. URLs: TCGA data matrix, https://tcga-data.nci.nih.gov/tcga/dataAccessMatrix.htm; TCGA Research Network, http://cancergenome.nih.gov/.

## Acknowledgements

We thank Peng Zhang, Ying Lu and Xuebing Guan for help with preparation and administration of the experimental materials, Dr. Yanzhong (Frankie) Yang from city of Hope providing the PRMT plasmids, and members of the Wei laboratory for their suggestions and discussions of this project. This work was supported by the National Natural Science Foundation of China (82073082; 82311530050; 81772948; 81972617), the Shanghai Interactional Collaborative Project (23410713300), the National Key Research and Development Program of China (2022YFC2703600), Key Research and Development Program of Anhui Province (2022i01020023), Jane ja Aatos Erkon säätiö, Sigrid Juséliuksen Säätiö, Syöpäjärjestöt, and Fudan University Recruit Funding. The organizations that offered the grants played no role in study design, data generation and analysis, decision to the preparation and publication of the manuscript. Linux High-Performance Computing servers were supported by the Medical Research Data Center in Shanghai Medical College of Fudan University and the CSC-IT CENTER FOR SCIENCE LTD.

## Author contributions

C.L., and C.Z. designed the project and analyzed most of the experiments. G.-H.W. conceived and designed the study, and provided supervision, suggestions and guidance. C.L., C.Z., and P.T. performed most of the experiments. C.L., Q.Z., and M.Z. performed the in vivo experiments. Qin Zhang, and Z.W. contributed to bioinformatics analysis. C.L. and C.Z. performed IP-MS assays. Q.T., Y.W., Y.Z., and Z.L. provided patient samples and contributed to TMA evaluation. L.Q.Z., M.L., X.K., H.Z., and Q.Y.L. provided resources and suggestions. J.C. contributed to structural biology analysis. C.L. and G.-H.W. wrote the manuscript with input from C.Z., P.T. and other responsible authors. All authors involved in the discussion of the results and the contributions to the manuscript.

## Declaration of interests

The authors declare no conflicts of interest.

## Notes

### Competing Interest Statement

The authors have declared no competing interest.

