## Supplemental Figure 1-8 for "Epigenetic reader ZMYND11 noncanonical function restricts HNRNPA1-mediated stress granule formation and oncogenic activity"

### Lian *et al.* Supplemental Figure 1

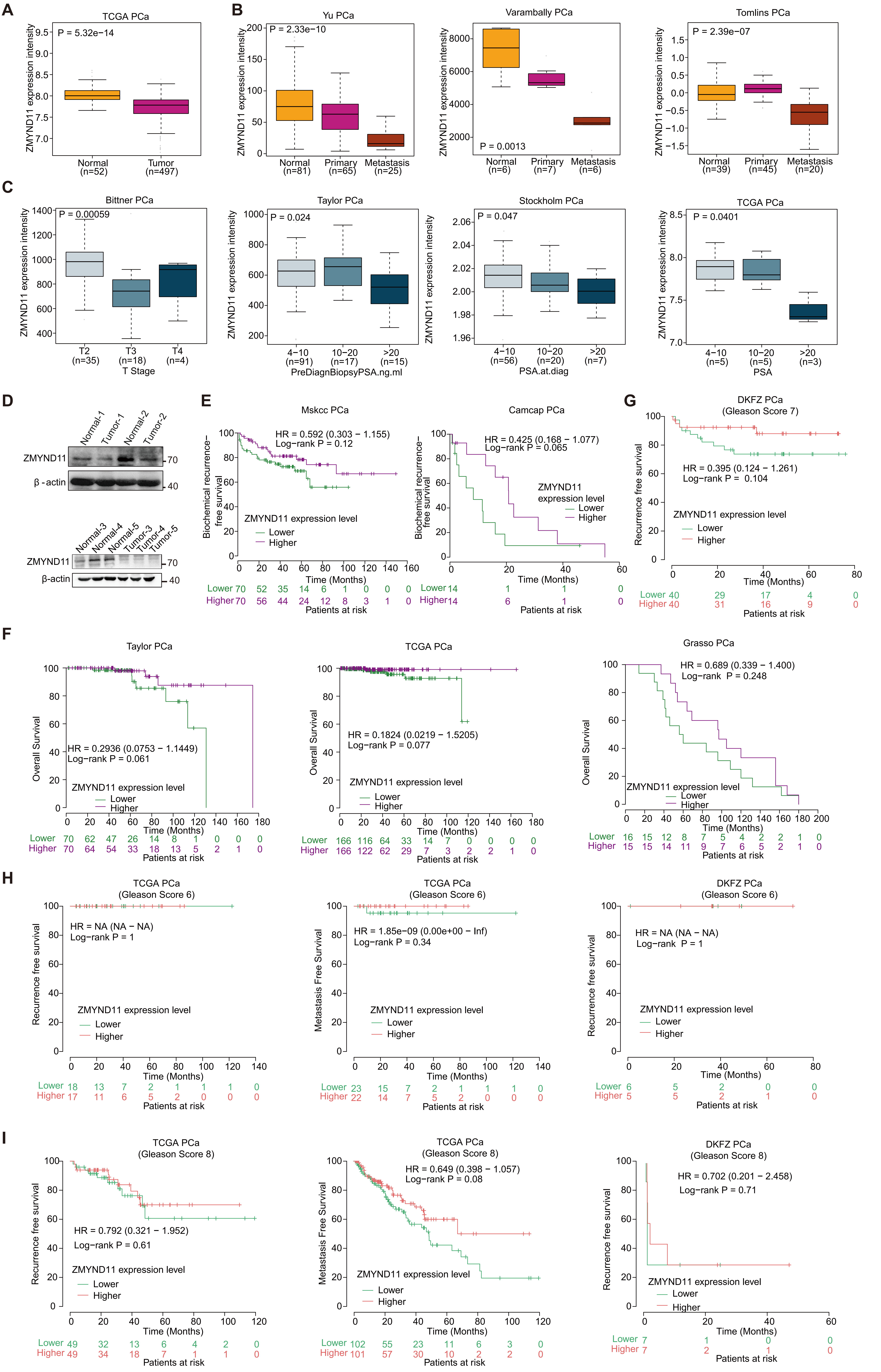

**Figure S1: ZMYND11 downregulation is prevalent and consistently correlates poor survival outcomes in patients with PCa.**

(A) ZMYND11 mRNA expression was significantly downregulated in human prostate tumors in TCGA PRAD dataset.

(B) ZMYND11 expression levels were significantly downregulated in human metastatic prostate tumors.

(C) Lowered ZMYND11 expression was greatly associated with high T stage and serum PSA levels in PCa.

(D) Western blotting of the ZMYND11 protein levels in five tumor-benign paired tissues of murine prostate.

(E) Kaplan-Meier analysis of biochemical recurrence-free survival in patients with prostate tumors expressing high or low levels of ZMYND11 in two independent cohorts of PCa.

(F) Kaplan-Meier analysis of overall-survival in patient groups with prostate tumors expressing high or low levels of ZMYND11 in three independent cohorts of PCa.

(G) Lower expression levels of ZMYND11 indicate predictive values for recurrence-free survival in patient group with Gleason score 7 (intermediate-risk PCa).

(H-I) Expression levels of ZMYND11 indicates no predictive values for recurrence and metastasis-free survival in patient group with Gleason score 6 (low-risk PCa) or Gleason score 8 (high-risk PCa). In E-I, the p values were examined by the log-rank tests.

#### Lian *et al.* Supplemental Figure 2

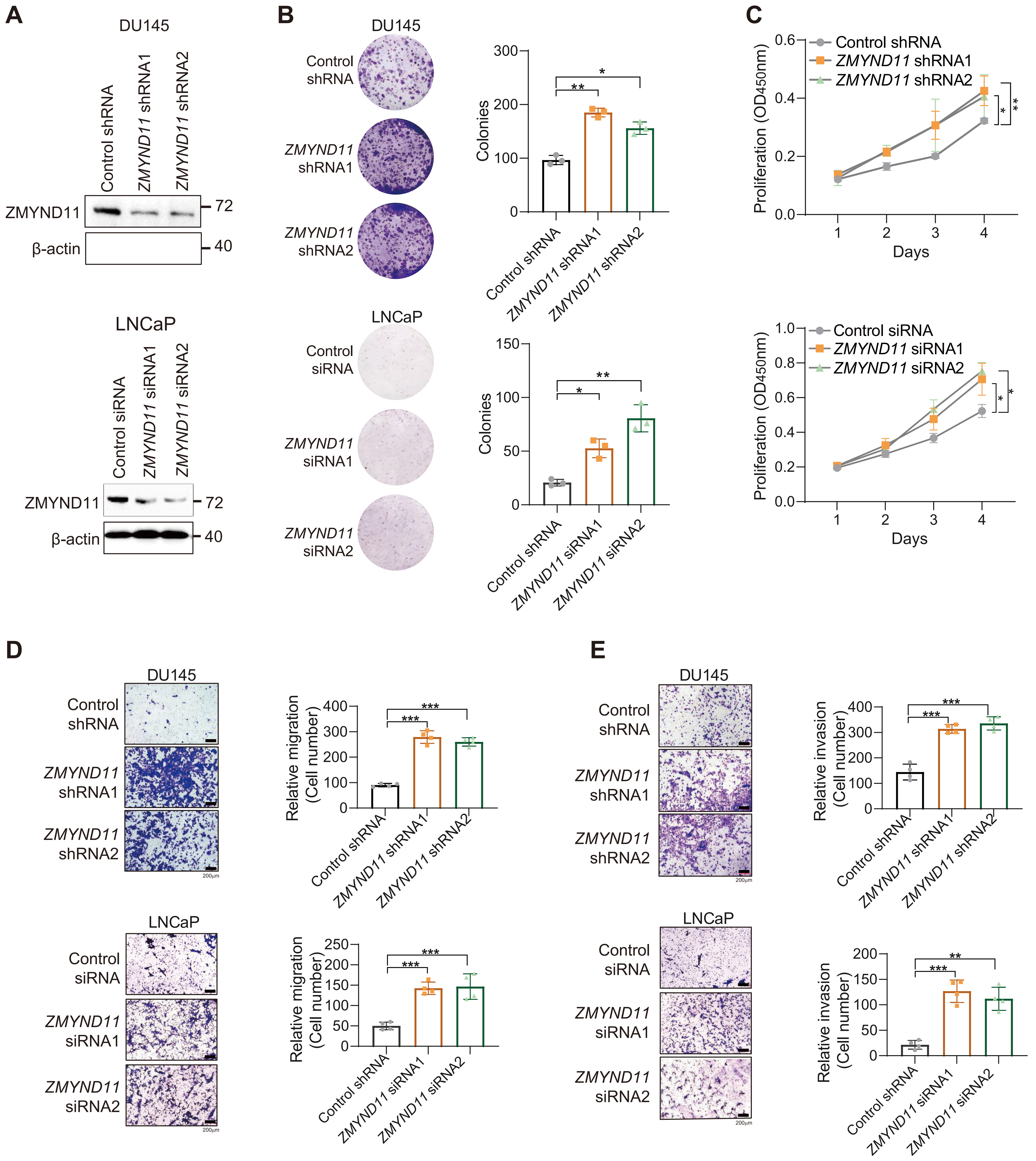


**Figure S2: Dampened ZMYND11 expression accelerates PCa cell proliferation and metastasis.**

(A) Efficiency of siRNA or shRNA-mediated knockdown of ZMYND11 was measured using western blot.
(B) Colony-forming units of DU145 and LNCaP cells transfected with control and shRNAs or siRNAs against ZMYND11, respectively.
(C) Proliferation rates of DU145 and LNCaP cells with ZMYND11 depletion or control were measured at the indicated time points (absorbance at 450 nm; mean ± s.d. of three independent experiments).
(D and E), Representative images and quantification of relative migration
(D) and invasion (E) for the cells transfected with control or shRNAs against ZMYND11. Scale bar, 200 μm. In B-E, data shown are mean ± SD of three technical replicates. *P<0.05, **P<0.01, ***P<0.001. P values were assessed by unpaired two-tailed Student's t-test.

#### Lian *et al.* Supplemental Figure 3

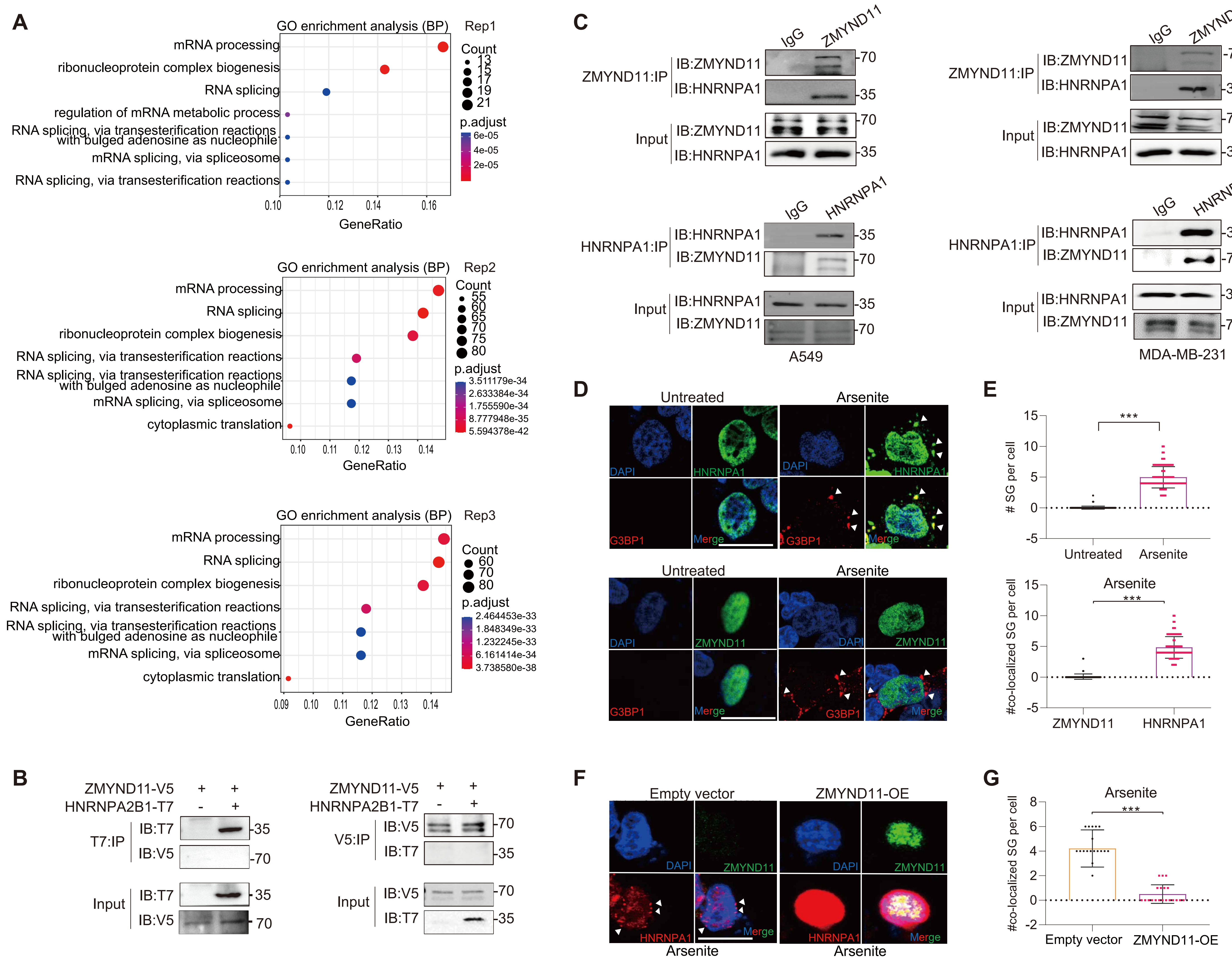


**Figure S3: ZMYND11 displays robust protein-protein interaction with HNRNPA1.**

(A) Gene Ontology (GO) annotation of proteins that identified in three replicates of mass spectrometry results for the ZMYND11 precipitated protein complexes. Top-enriched pathways are indicated.
(B) Immunoprecipitation (IP)-Western blot for analyzing the interaction between ZMYND11 and HNRNPA2B1 from HEK293T cells co-transfected with V5-tagged ZMYND11 and T7-tagged HNRNPA2B1. IB, immunoblot.
(C) Immunoprecipitation-Western blot for analyzing the endogenous interaction of ZMYND11 with HNRNPA1 in human lung cancer A549 and breast adenocarcinoma MDA-MB-231 cells. IgG, immunoglobulin. IB, immunoblot.
(D and E) Immunofluorescence colocalization analysis of ZMYND11, HNRNPA1 and G3BP1 in 22Rv1 cells treated with sodium arsenite (SA, 0.5mM) or control for 1h. Scale bar, 25 μm (D). SG numbers per cell and ZMYND11 or HNRNPA1 co-localized SGs per cell were quantified (E). Error bars, SD, n = 3 biological replicates. *p < 0.05, **p < 0.01, ***p < 0.001, Student's t-tests.
(F and G) 22Rv1 cells were co-transfected with HNRNPA1-T7 and ZMYND11-V5 or empty vector, then treated with sodium arsenite (SA, 0.5mM) for 1h followed by immunostaining analysis of stress granule (SG) formation (F). SGs per cell were quantified (G). Scale bar, 25 μm. *** p < 0.001, unpaired two-tailed Student's t-tests.

**Lian *et al.* Supplemental Figure 4**

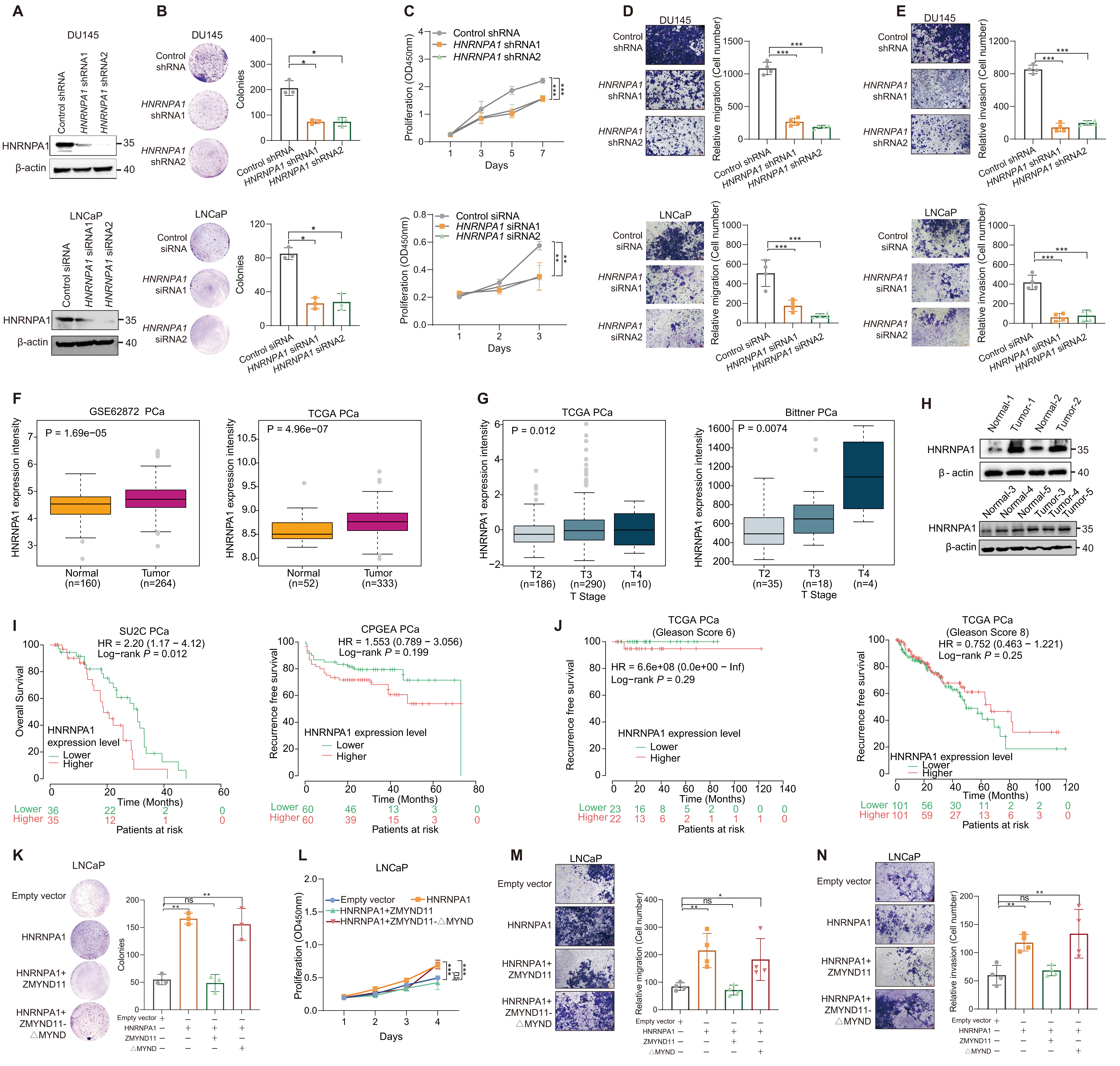

**Figure S4: ZMYND11 restricts the HNRNPA1 oncogenic function in an MYND domain dependent manner.**
(A) Western blot analysis of knockdown efficiency mediated by HNRNPA1-specific shRNAs or siRNAs in DU145 and LNCaP cells, respectively.
(B) Colony-forming units of DU145 or LNCaP cells transfected with control or shRNAs/siRNAs against HNRNPA1.
(C) Cell proliferation was measured at the indicated time points by Cell Counting Kit-8 (CCK-8) colorimetric assay (absorbance at 450 nm; mean ± s.d. of three independent experiments).
(D and E) Representative images and quantification of relative migration (D) and invasion (E) for the cells transfected with control and the shRNA or siRNA against HNRNPA1, respectively. Data shown are mean ± SD of three technical replicates. *P<0.05, **P<0.01, ***P<0.001, assessed by unpaired two-tailed Student's t-test.
(F) HNRNPA1 expression levels are significantly upregulated in human prostate tumors compared to benign specimens.
(G) HNRNPA1 upregulation shows significant association with high-tumor stage cancer in two independent patient cohorts of PCa.
(H) Protein levels of HNRNPA1 in five tumor-benign paired tissues of murine prostate.
(I) Kaplan-Meier overall-survival analysis in patient groups with prostate tumors expressing high or low levels of HNRNPA1 in two independent cohorts of human PCa patients.
(J) Kaplan-Meier analysis showed that HNRNPA1 indicated poor predictive values for recurrence-free survival in patient groups with Gleason score 6 (low-risk PCa) and Gleason score 8 (high-risk PCa), respectively.
(K-N) Representative images and quantification of colony-formation (K), cell proliferation (L), migration (M), and invasion (N) for LNCaP cells transfected with empty vector, HNRNPA1 alone, or together with the indicated full-length ZMYND11 or MYND-domain deletion mutant. Mean ± SD, p value was assessed using two-tailed Student's t-test. *P<0.05, **P<0.01, ***P<0.001. Scale bars, 25 μm.

**Lian *et al.* Supplemental Figure 5**

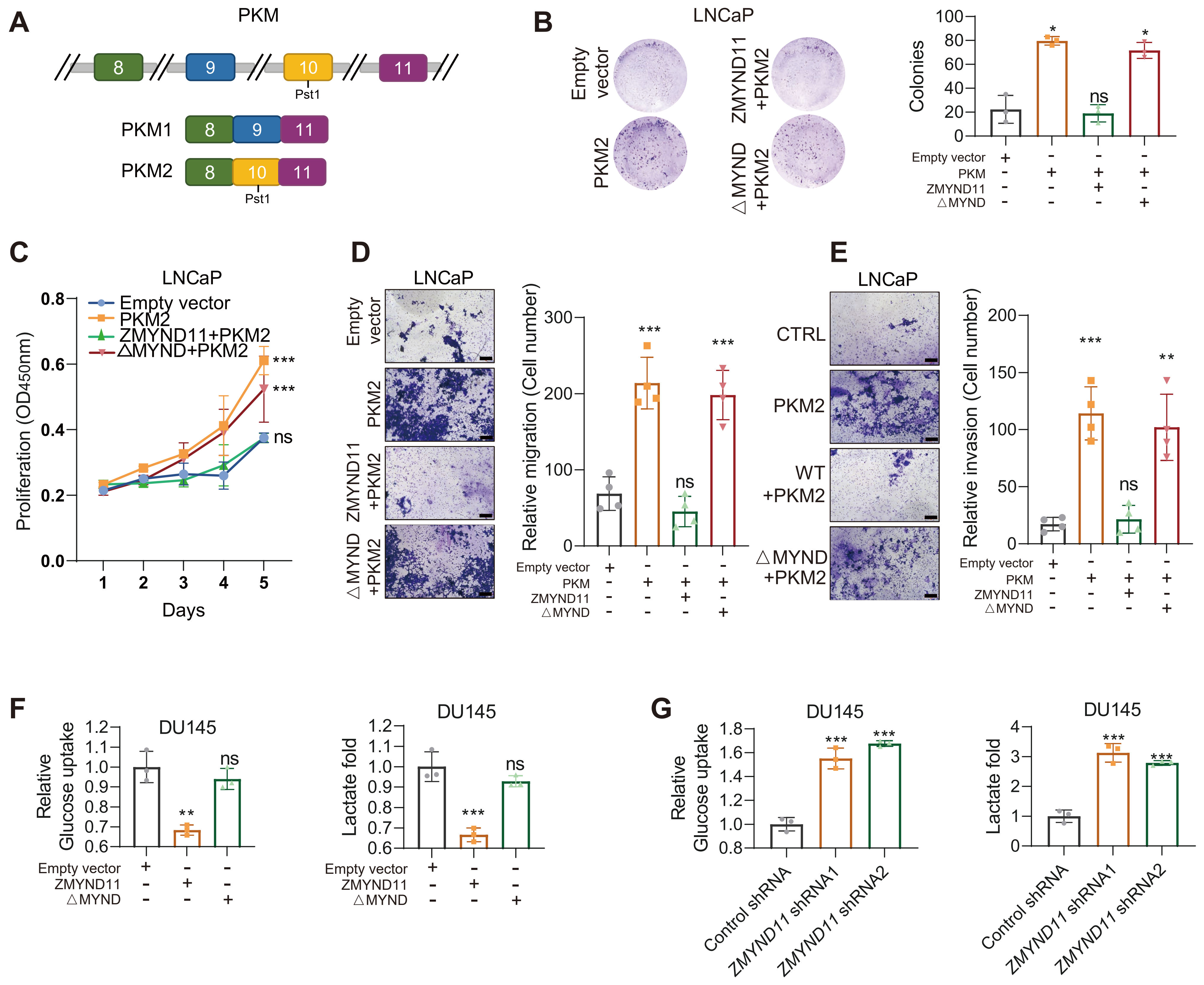

**Figure S5: ZMYND11 regulates HNRNPA1-mediated alternative splicing of PKM and counteracts PKM2-induced aggressive cellular phenotype in PCa.**

(A) Scheme for PKM1/PKM2 transcript forms and assaying PKM1/PKM2 ratios in human cells.

(B-E) Representative images and quantification of colony formation (B), cell proliferation (C), migration (D) and invasion (E) in LNCaP cells co-transfected with PKM2 alone or together with either the full-length ZMYND11 or the MYND-domain-deletion mutant. Error bars, SD (B-E). ns, not significant, *p < 0.05, **p < 0.01, ***p < 0.001, two-tailed Student's t tests.

(F-G) Relative glucose uptake and lactate production in DU145 cells overexpressing ZMYND11 or its MYND-domain-deletion mutant (F) and stably expressing ZMYND11 shRNA (G) compared to control cells. ** p< 0.01, *** p < 0.001; ns, not significant, assessed by the unpaired two-tailed Student's t-tests.

#### Lian *et al.* Supplemental Figure 6

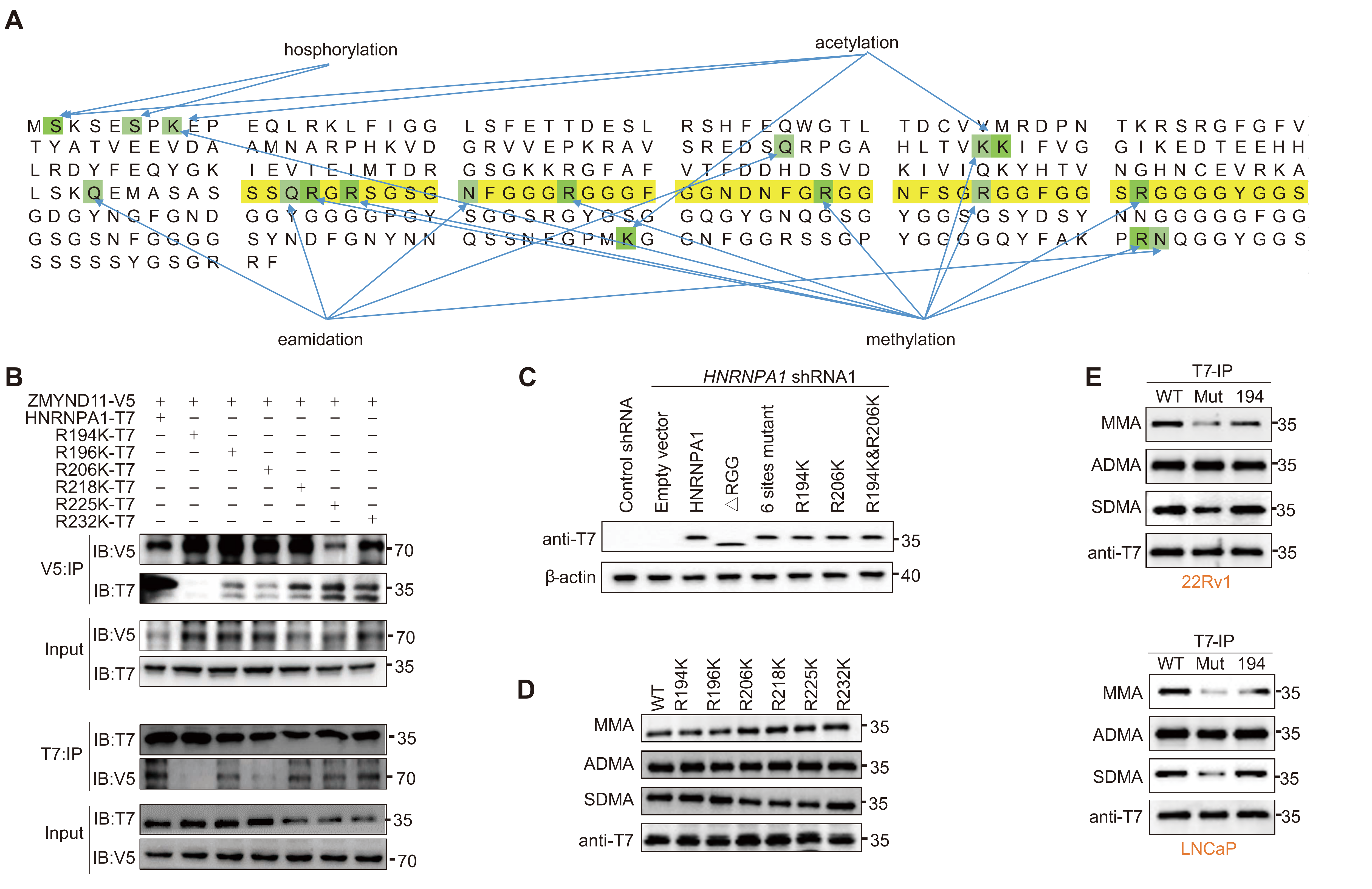


**Figure S6: HNRNPA1 arginine 194 methylation is required for its oncogenic function and recognition by ZMYND11.**

(A) Mass-spectrometry analysis of HNRNPA1 post-translational modification. RGG domain residues are shown in yellow. Amino-acid residues with potential post-translational modifications (PTM) are indicated in green.
(B) The lysates from HEK293T cells co-expressing V5-tagged ZMYND11 and T7-tagged wild type HNRNPA1 or the indicated mutant constructs were subject to immunoprecipitation with anti-V5 antibody followed by immunoblotting (IB). The reciprocal co-IP results were shown in the bottom.
(C) Western blot showed ectopic expression of different plasmids in 22Rv1 cells with the shRNA-mediated knockdown of HNRNPA1.
(D) The lysates from 22Rv1 cells expressing T7-tagged wild type (WT) HNRNPA1 or the indicated mutants were subject to immunoprecipitation using anti-T7 antibody followed by immunoblotting (IB). Three types of arginine methylation were also analyzed.
(E) Ectopic expression of HNRNPA1 or its variants was subject to immunoprecipitation with anti-T7 antibody and three types of arginine methylations were analyzed in the indicated cell types. 22Rv1 and LNCaP cells were transfected with T7-tagged WT HNRNPA1 or the indicated mutants either with complete mutation of six arginine amino residues or R194 un-mutant alone.

#### Lian *et al.* Supplemental Figure 7

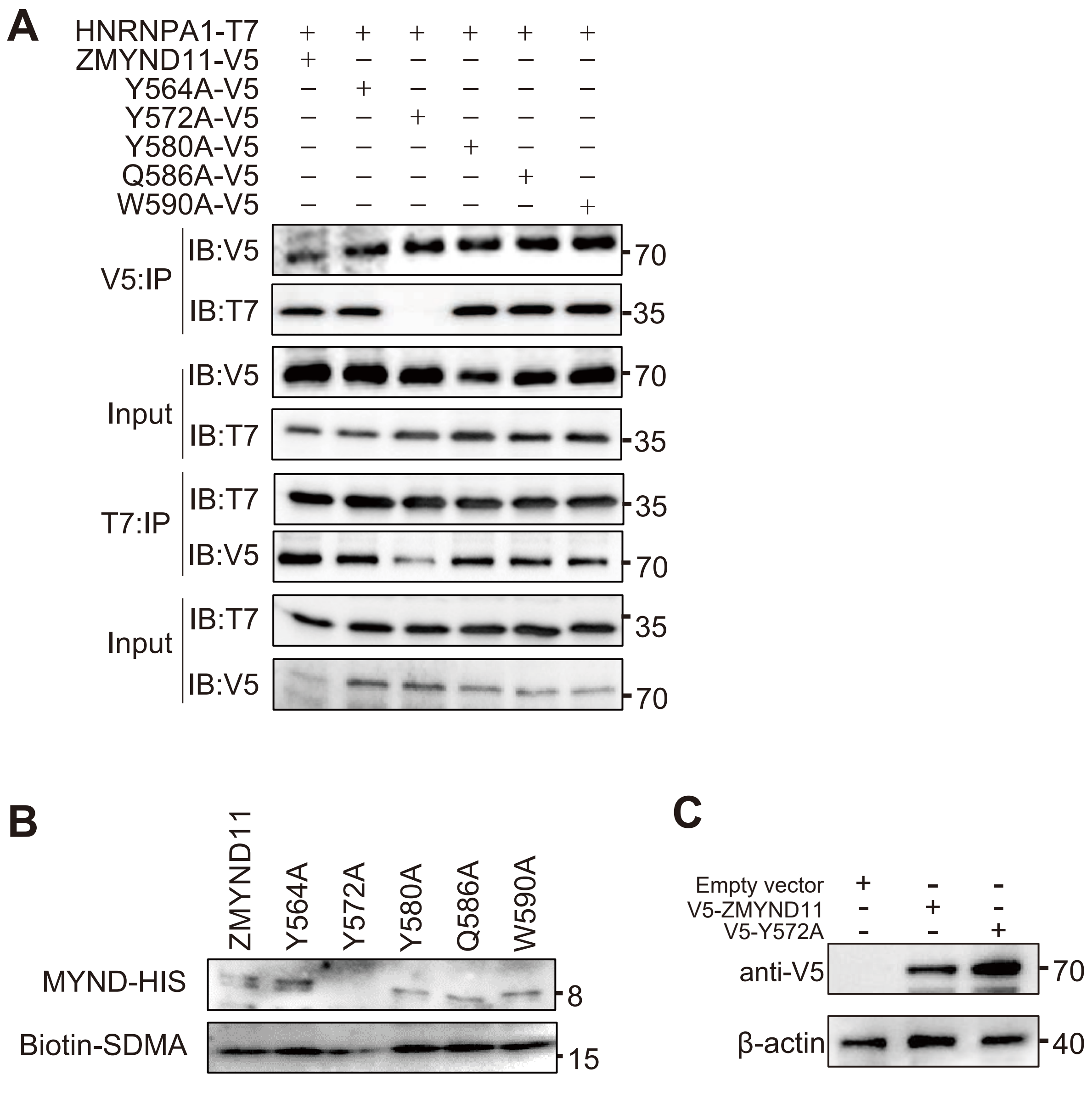


**Figure S7: Tyrosine residue 572 of ZMYND11 is functionally important and responsible for specifically reading SDMA on HNRNPA1.**

(A) The lysates from HEK293T cells expressing T7-tagged HNRNPA1 and V5-tagged ZMYND11 or the mutants: Y564A, Y572A, Y580A, Q586A and W590A, were immunoprecipitated with anti-V5 and anti-T7 antibody, respectively, followed by immunoblotting (IB).

(B) Pull-down assays of the interaction between HIS-tagged MYND domain of ZMYND11 and the mutants and the biotinylated peptide containing R194 SDMA.

(C) Western blot detected the protein extracts of 22Rv1 cells transfected with wild type (WT) ZMYND11 or the Y572A mutant.

#### Lian *et al.* Supplemental Figure 8

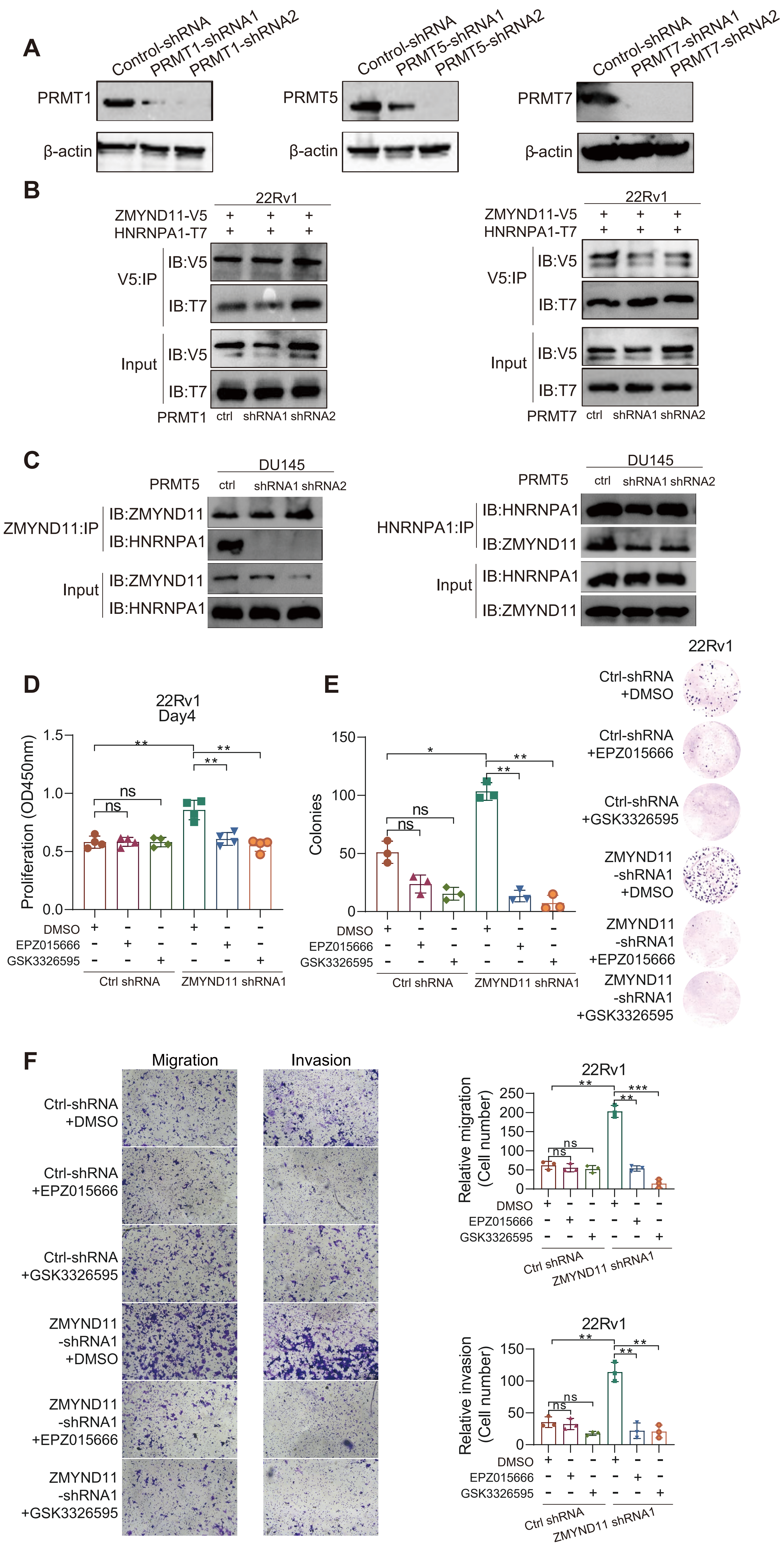

**Figure S8: Pharmacological inhibition of PRMT5 impairs physical association of ZMYND11-HNRNPA1 and attenuates the growth of ZMYND11-low expression PCa cells.**

(A) Western blot examination of the stable knockdown of PRMT1, PRMT5 or PRMT7 in HEK293T cells.
(B) The lysates from 22Rv1 cells stably expressing control or shRNAs against PRMT1 or PRMT7 were subject to co-expression of V5-tagged ZMYND11 and T7-tagged HNRNPA1 followed by co-immunoprecipitation-immunoblotting (IB) with anti-V5 antibody.
(C) Immunoprecipitation-immunoblotting (IB) analysis of an endogenous interaction between ZMYND11 and HNRNPA1 in DU145 cells stably transfected with control and ZMYND11-targeting shRNAs.(D-F) Representative images and quantification of cell proliferation (D), colony formation (E), migration and invasion (F) of 22Rv1 cells transfected with ZMYND11-targeting shRNAs and treated with PRMT5 inhibitors (10μM, 48h). Error bars, SD (D-F). *p < 0.05, **p < 0.01, ***p < 0.001, two-tailed Student's t tests.
